# Modulating the epigenetic state promotes the reprogramming of transformed cells to pluripotency in a line-specific manner

**DOI:** 10.1101/2022.12.01.518778

**Authors:** Xiuling Fu, Qiang Zhuang, Isaac A. Babarinde, Liyang Shi, Gang Ma, Haoqing Hu, Yuhao Li, Jiao Chen, Zhen Xiao, Boping Deng, Li Sun, Ralf Jauch, Andrew P. Hutchins

## Abstract

Somatic cell reprogramming and oncogenic transformation share surprisingly similar features, yet transformed cells are highly resistant to reprogramming. There must be barriers that block transformed cells from reprogramming, but the nature of those barriers is unclear. In this study, we generated a systematic panel of transformed mouse embryonic fibroblasts (MEFs) using a variety of oncogenic transgenes, and discovered transformed cell lines that remain compatible with reprogramming when transfected with *Oct4*/*Sox2*/*Klf4*/*Myc*. By comparing the reprogramming-capable and incapable transformed lines we identified multiple stages of failure in the reprogramming process. Some transformed lines failed very early, whilst other lines seemed to progress through a normal-looking reprogramming process. Finally, we show that MEK inhibition overcomes one critical reprogramming barrier by indirectly suppressing a hyperactive epigenetic state in some of the transformed cells. This study reveals that the barriers underlying resistance to reprogramming vary between the different transformation methods.

**Key findings:**

- Somatic cell reprogramming of transformed cells is context-specific
- Inhibition of MEK converts some cell lines to reprogramming-capable
- Transformed cell lines are characterized by a hyperactive chromatin state
- MEK inhibition indirectly affects chromatin to enable reprogramming

## Introduction

Transformed cells and embryonic stem cells (ESCs) have a remarkable list of similarities [1]. Both cell types have a relaxed chromatin state [2], adopt a glycolysis-biased metabolism despite the availability of oxygen [3, 4], can undergo an epithelial-mesenchymal transition (EMT) [5], and form teratomas when injected into immunocompromised mice [6]. Transformed cells acquire features reminiscent of embryonic development, such as increased cellular plasticity and the up-regulated expression of pluripotent genes, including *OCT4, NANOG*, and *SOX2* [7]. Indeed, expression of pluripotent-specific genes in patient tumor samples correlate with poor clinical outcomes [8, 9]. For example, overexpression of *Oct4* led to epithelial lesions [10], and *NANOG* expression is associated with colorectal [11] and prostate cancer [12].

Somatic cells can be reprogrammed to induced pluripotent stem cells (iPSCs) by the transfection of a cocktail of transgenes, particularly *Oct4* (*Pou5f1*), *Sox2, Klf4*, and *Myc* [13]. Curiously, despite being an artificial process, the reprogramming of somatic cells to iPSCs passes through distinct phases, reminiscent of a developmental program [14-16]. Tumorigenic transformation also passes through a series of distinct phases in a process that has similarities to somatic cell reprogramming [1, 17]. Some studies suggest direct links between reprogramming and cancer. For example, transient *in vivo* activation of reprogramming factors leads to tumor formation [18, 19], and cancer-associated mutations in the transcription factor *SOX17* can confer reprogramming capability to the normally incapable SOX17 [20].

There have been reports on reprogramming primary human cancer cells to an embryonic state, including cancerous cells from liver, gastrointestinal tract, and sarcoma cell lines [21-24]. Additionally, blood cancer cells seem more amenable to be reprogrammed: there are reports on reprogrammed cells generated from human KBM7 cells, T cell acute lymphoblastic leukemia, acute myeloid leukemia [25-27], lymphoblastoid cells [28, 29] and chronic myeloid leukemia cells [24, 30-32]. However, these blood cells seem to be the exception, since in the general transformed cells are resistant to reprogramming [33]. Importantly, it is unclear how close those iPSC-like cells are to genuine iPSCs, and the iPSC-like cells may not be completely reprogrammed. For example, whilst the overall gene expression of the reprogramed cells is closer to iPSCs than the parental lines, it remains distinct from genuine ESCs/iPSCs [23, 28, 34]. It is also possible that reprogramming from primary tissues is selecting for bystander untransformed cells [35]. Finally, even when reprogramming appears possible, the efficiency is much lower than the already low efficiency of wild-type reprogramming. This is a curious contradiction. Considering the similarities between cancer cells and iPSCs and the pathways used to generate them, it seems that reprogramming should be easier in transformed cells as cancer cells often have reduced barriers to cell type conversion.

To explore the barriers blocking the reprogramming of transformed cells, we generated a panel of eight artificial transformed mouse cell lines. Seven of these lines can acquire OCT4-GFP+ reporter expression and pluripotent characteristics using a conventional OSKM-reprogramming protocol. Albeit at very low efficiency. Of the remaining three reprogramming-incapable lines, we show that the defects in reprogramming are line-specific. Reprogramming is a phased process and some lines fail in the early phases, others in the later phases. Compared to wild-type cells, the reprogramming-incapable lines show a heightened ‘hyperactive’ chromatin state and demonstrate global increases in chromatin accessibility and histone acetylation. Some of the transformed lines could be converted to be reprogramming-capable by inhibiting MEK signaling which moderates the active chromatin state and leads to decreased histone acetylation.

## Results

### Mouse transformed cell lines are recalcitrant to reprogramming

To explore reprogramming in transformed cell lines we first attempted to reprogram mouse cell lines that were originally derived from a tumor or that were spontaneously transformed. These cell lines have been grown in culture for an extended period and their passage number is unknown. We chose three mouse cell lines: 3T3-L1, a spontaneously immortalized fibroblastic cell line; 4T1, a metastatic breast tumor cell line; and N2a (Neuro-2a), a neuroblastoma cell line (**Figure 1a**). We attempted to reprogram these cell lines using OSKM in serum+Vc conditions [36]. For the 3T3-L1 and 4T1 (but not the N2a), we observed colonies that morphologically resembled iPSCs on day 15 (**Figure 1a**). Low levels of NANOG protein could be detected by immunofluorescence in some 3T3-L1 and 4T1 colonies (**Figure 1b**). We attempted to manually pick the iPSC-like colonies and establish iPSC lines. However, contaminating 3T3-L1 and 4T1 cells would outcompete any iPSC-like colonies, and based on morphology, the cultures would rapidly revert to homogenous 3T3-L1 or 4T1 cultures. Hence, we sorted reprogrammed cells positive for ICAM and negative for CD44 (ICAM+/CD44-), which marks the late stage of reprogramming in wild-type MEFs [37]. ICAM+/CD44-cells were sorted from the reprogrammed 3T3-L1 and 4T1 cells, and we detected small numbers of ICAM+/CD44-cells at D15 of reprogramming in 3T3-L1 and 4T1, but not N2a (**Figure 1c, d**). qRT-PCR of the ICAM+/CD44-cells indicated that *Esrrb* and *Nanog* were up-regulated in the 4T1, and *Essrb, Nanog*, and endogenous-*Pou5f1* in 3T3-L1 (**Figure 1e**). However, when the sorted cells were replated they rapidly reverted to the original cell type morphology and there was no evidence of iPSC-like cells (**Figure 1f**). Overall, these results highlight the difficulty in reprogramming transformed cell lines to iPSCs.

**Figure 1.**
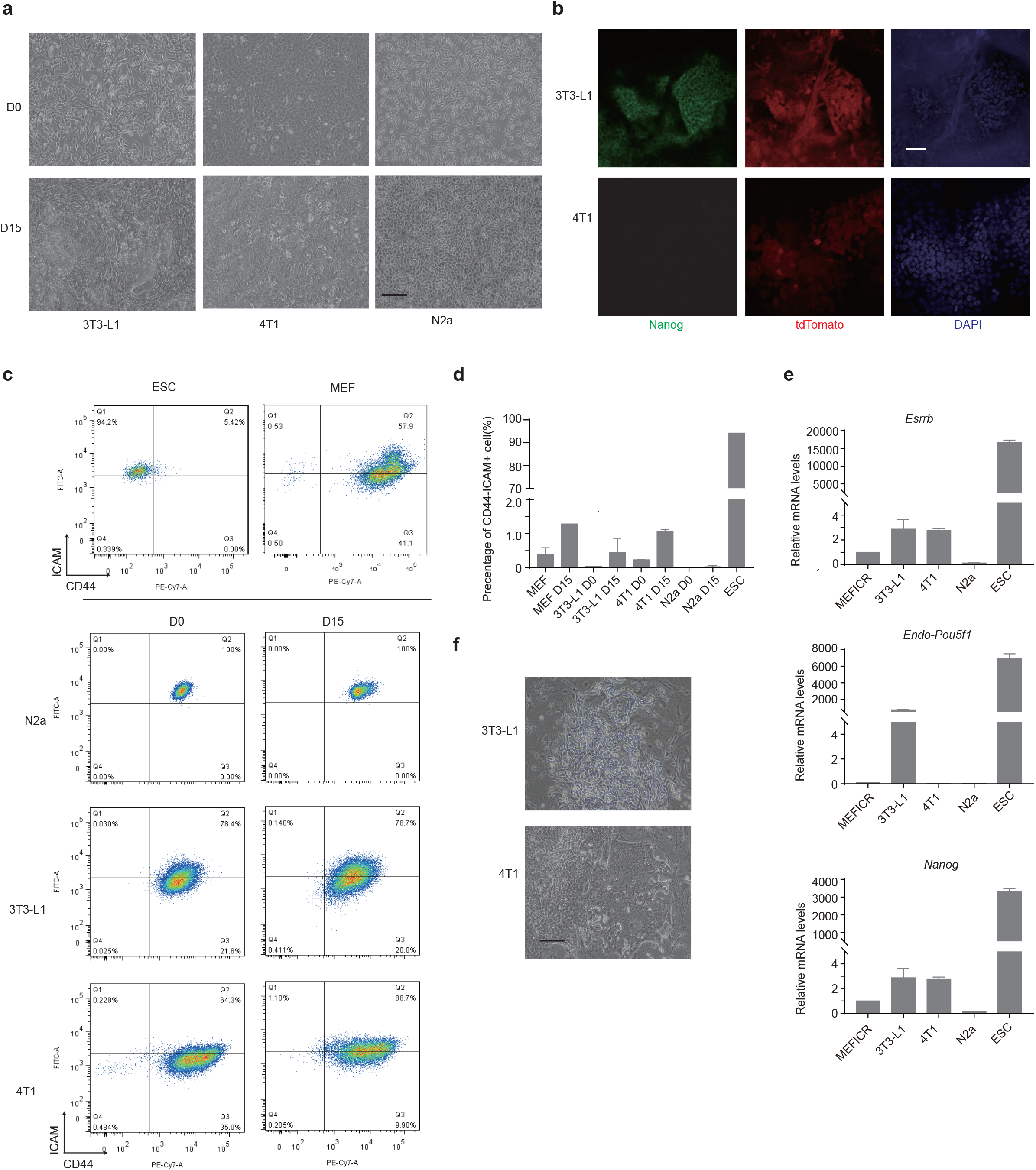
Transformed cell lines are highly resistant to reprogramming. **a** Bright-field images of the reprogramming of 3T3-L1, 4T1, and N2a cell lines in serum+Vc reprogramming conditions. Images are shown from day 0 (D0) or day 15 (D15) of the reprogramming experiment. Morphologically iPSC-like cells can typically be observed at D15 and onwards in wild-type reprogramming. Scale bar=100 *µ*m. **b** Fluorescent microscopy images for NANOG and tdTomato expressed from the OSKM transgene cassette for the reprogramming of 3T3-L1 and 4T1 cells. Scale bar=100 *µ*m. **c** FACS (fluorescent activated cell sorting) plots for wildtype MEFs and mESCs and cells from a reprogramming time course at day 0 or 15 from 3T3-L1, 4T1, and N2a cells. **d** Bar chart of the percentage of ICAM+/CD44-sorted cells in the indicated cell lines at day 0 or day 15 of reprogramming. Data is from the mean of three biological replicates. **e** RT-qPCR showed the indicated gene expression from pooled cultures of the sorted ICAM+ and CD44-cells from the indicated cell lines five days after replating. The mean of two biological replicates ± s.d. is shown. **f** Bright-field images of the sorted ICAM+ and CD44-cells from the indicated cell lines five days after replating. Scale bar=100 *µ*m.

### Features of untransformed MEF senescence in relation to reprogramming

We first explored wildtype MEF reprogramming to understand the blocks on reprogramming in the normal case so that we can rule them out in the transformed cell lines. Wildtype MEFs can only reprogram at early passages [38], and the efficiency declines rapidly before complete failure after passage 4 (**Figure S1a**), when using a conventional reprogramming OSKM (*Oct4, Sox2, Klf4*, and *Myc*) system [36]. RNA-seq of MEFs from passages 1-6 revealed changes in cell cycle genes, including the downregulation of cyclins and other positive cell cycle regulators and the activation of cell cycle inhibitors, particularly *Cdkn1a* (**Figure S1b**). We divided the samples into MEFs that could reprogram successfully (P1, 2) or failed (P5, 6) and measured significantly differentially expressed (DE) genes. In total, 337 genes were significantly up-regulated and 419 significantly down-regulated (**Figure S1c, d**). Gene ontology (GO) and gene set enrichment analysis (GSEA) of the differentially expressed genes indicated that the down-regulated genes were related to the cell cycle and extracellular matrix. Up-regulated genes were related to cell migration/adhesion, MAPK activity, apoptosis, and inflammation (**Figure S1e, f**), possibly caused by reactive oxygen species (**Figure S1f**). The inflammatory response is exemplified by the up-regulation of cytokines and chemokines in the P5,6 MEFs (**Figure S1g**). We defined the up-regulated gene set as the ‘MEF senescent signature’ and the down-regulated gene set as a ‘reprogramming permissive signature’ (**Supplementary Table S1**).

### Generation of a panel of oncogenic transformed MEF cell lines

As the above-transformed cell lines have been maintained in culture for an extended time they have likely accumulated many genetic alterations to adapt to the cell culture environment. These changes may permanently impair their ability to reprogram. Consequently, to achieve a controlled system we generated immortalized MEFs using 9 different combinations of factors to represent a spectrum of transformation. The factors were chosen to cover different methods of immortalization: oncogenic transcription factors (*Myc*, Hras^G12V^, Mef2d, p53DD), viral transforming factors (SV40T, E1A), apoptotic factors (Bcl2), and an engineered epigenetic factor (Hdac7SA; serines at positions 178, 344, and 479 substituted with alanine, to block nuclear export) [39-41]. The sequences of the transformation factors are in **Supplementary Table S2**. Not all of these factors could successfully transform MEFs, and, except for SV40T, at least two factors were required for successful transformation (**Figure S2a, b**). The immortalized cell lines were maintained for at least 1 month to remove any non-transformed background MEFs. After 1 month resulting lines still expressed the appropriate transgenes, as confirmed by RNA-seq, RT-qPCR, and Western blot (**Figure S2c-e**). The exception was the p53DD.Myc lines, as the p53DD transgene was silenced at the protein level, whilst *Myc* could be detected by both RT-qPCR and Western blot (**Figure S2c-e**). This suggests that both transgenes are required to generate immortal MEFs, but once immortalized, p53DD becomes dispensable.

We next looked at the features of the transformed cells. All of the immortalized MEFs proliferated faster than the wild-type MEFs (**Figure S2f**). We tested five lines for the capability to form tumors in nude mice, Hras.E1A, SV40T, and Hras.Myc could form tumors, whilst Bcl2.Myc and Hdac7SA.Mef2d could not (**Figure S2g, h**). Hoechst staining of cross sections through the tumors indicated various differentiation layers, although mainly mesoderm (**Figure S2i**). Aneuploidy is a common feature of cancer, although the role of aneuploidy in transformation remains unclear [42]. A normal karyotype is required for post-implantation embryonic development [43], but may not be incompatible with pluripotency [44-46]. To rule out the impact of karyotype abnormalities on reprogramming capability we confirmed a normal karyotype for four of the lines that the subsequent study will mainly focus on (**Figure S2j**).

### Somatic cell reprogramming reveals a spectrum of reprogramming capability

Using this panel of immortalized MEF cell lines we reprogrammed them using a lentiviral OSKM system with vitamin C, which is known to promote reprogramming [36]. We used transformed lines derived from OG2 MEFs, which contain an Oct4-GFP reporter [36]. Surprisingly, several lines could give rise to GFP+ colonies. We labeled the lines as ‘succeeding’ (Hdac7SA.Mef2d, p53DD.Myc), ‘struggling’ (Bcl2.Myc, Hdac7SA.E1A, p53DD.E1A, Hdac7SA.Myc), and ‘failing’ (Bcl2.E1A, Hras.E1A, Hras.Myc, and SV40T) (**Figure 2a, b**). It should be noted that reprogramming the transformed lines was less efficient than wild-type MEFs, except for the p53DD.Myc line. Complete reprogramming requires a passage before the pluripotency gene expression program can be stably established. This process partially relies on the fast-dividing iPSCs outcompeting the slow-growing/senescent wildtype MEFs, however, the transformed MEFs were also highly proliferative (**Figure S2f**) and would compete with the reprogrammed iPSCs. Hence, we FACS sorted the GFP+ cells and passaged them to establish iPSC lines in the absence of transformed MEFs. Pluripotency marker genes were robustly expressed only in the sorted cells (**Figure 2c**), and we validated the expression of these genes by RT-qPCR and Western blot (**Figure S3a, b**). Comparison of the RNA-seq to a panel of pluripotency genes and somatic genes indicated that the gene expression levels were close to ESCs (**Figure 2c**), and cross-correlation and principal component analysis (PCA) indicated the cells were similar to ESCs (**Figure 2d, e**). DPre was used to compare the gene expression in the GFP+ sorted lines to a panel of cell types [47, 48], and identified most lines as ESC-like (**Figure S3c, d**). DPre also suggested that some ESC lines (e.g. Hdac7SA.Mef2d GFP+ line #4) were a mixture of ESCs and contaminating immortalized MEFs (**Figure S3d**). We confirmed that the iPSC lines could form teratomas with tissues representing all three germ layers (**Figure S3e**). A mark of complete reprogramming is the silencing of the OSKM transgenes, and the OSKM transgene was not detected in RNA-seq data (**Figure S3f**). Interestingly, the immortalization factors were only partially silenced. All were silenced except E1A in the Bcl2.E1A GFP+ line and Myc in the Bcl2.Myc GFP+ lines (**Figure S3f**). We confirmed Hdac7SA and Mef2d were silenced by qRT-PCR (**Figure S3g**). Overall, these data confirmed the successful generation of iPSC-like lines from some transformed tumorigenic normal karyotype MEF lines.

### Transformed MEFs have starting problems that impact reprogramming

We next explored the properties that made some immortal MEFs reprogramming-capable and others –incapable. We focused on two (non-exclusive) models: the inability to reprogram occurs due to problems in the originating MEFs, or it is caused by failures to traverse the correct sequence of events required for successful reprogramming.

**Figure 2.**
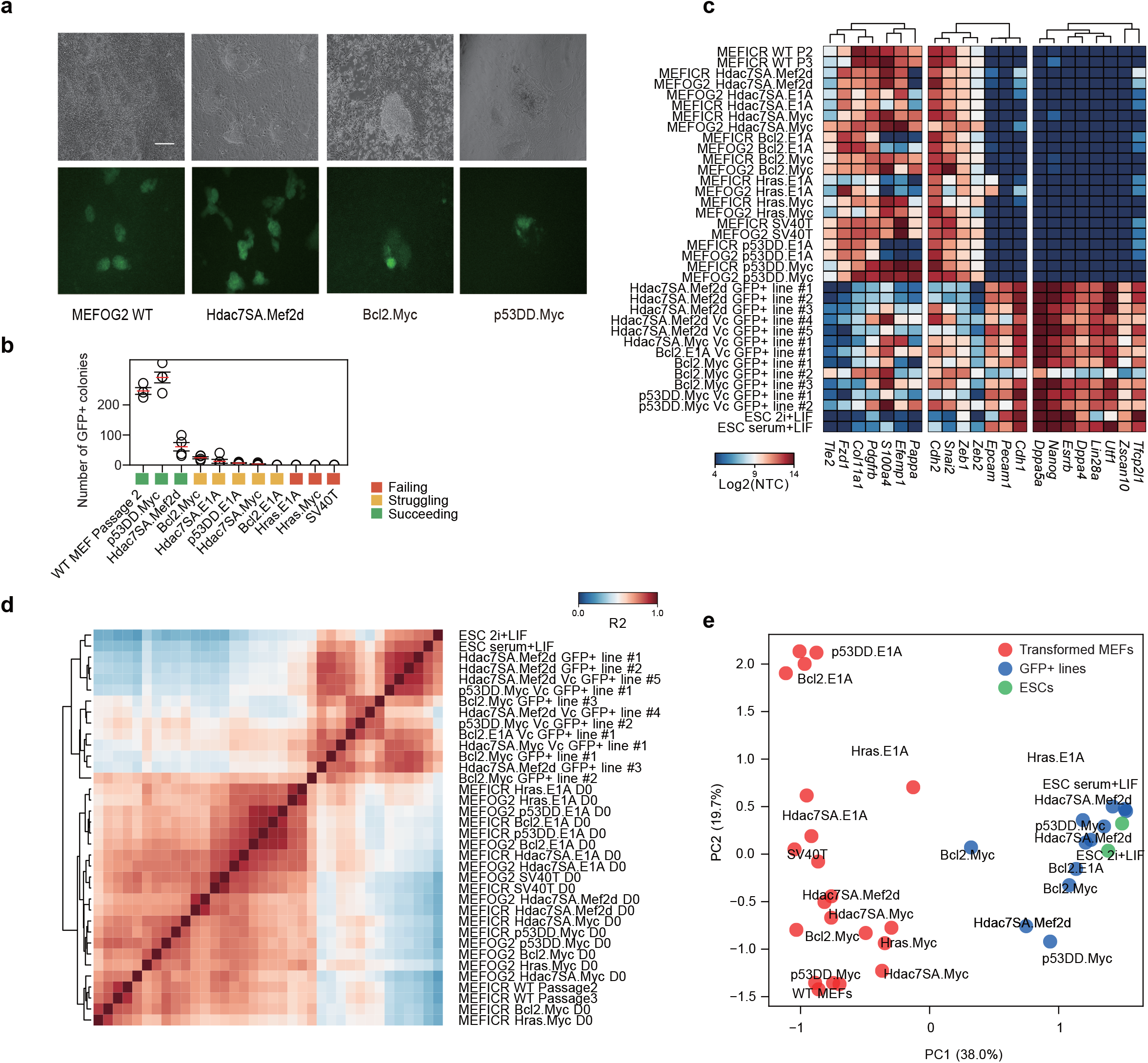
Characterization of transformed fibroblast cell lines capable and incapable of reprogramming to iPSC-like cells. **a** Representative brightfield and GFP images from reprogrammed transformed MEFs for the indicated cell lines. Scale bar=100 *µ*m. **b** Chart showing the number of Oct4-GFP+ colonies from the indicated transformed MEF lines. Each MEF line was at least 1 month old, the experiment was performed at least three times for each cell line. Circles represent biological replicates; the red line indicates the mean and the error bars indicate the standard error of the mean (SEM). **c** Heatmap from the RNA-seq data showing the expression of a representative set of genes specific to somatic cells, pluripotent stem cells, or from the MET (mesenchymal-epithelial transition). NTC=normalized tag count. **d** Correlation heatmap (R^2^) of the original MEF samples and the reprogrammed GFP+ cells and ESCs. **e** PCA showing the first two dimensions of the RNA-seq data from the original wildtype or transformed MEF lines (red) and the GFP+ cell lines (blue) and ESCs (green) maintained in serum+LIF or 2i+LIF.

Immortalization of somatic adult cells occurs by a range of mechanisms and is accompanied by changes in the epigenetic state and gene expression patterns. We analyzed the gene expression of the starting MEF cells and observed a wide range of gene expression changes, with some MEF lines adopting substantial changes in gene expression, whilst other cell lines were less affected (**Figure 3a**). There was no simple correlation between the number of gene changes and reprogramming capability (**Figure 3a**). Despite a large number of gene expression changes, there was no evidence that the MEFs were transdifferentiating as their global gene expression patterns continued to correlate well against wildtype MEFs (**Figure 3b**), and DPre continued to identify the cells as MEF-like (**Figure S4a**). Gene ontology of the DE genes suggested changes were associated with basic cellular processes such as up-regulation of metabolic processes, and extracellular matrix genes, and down-regulation of ribosomes, cell cycle, and DNA repair processes (**Figure 3c, S4b**), which is the opposite of the senescent-like phenotype up-regulated in human arrested embryos [49]. Indeed, the MEF-senescent gene signature (defined in **Figure S1c**) was not up-regulated in the transformed lines, and the majority of lines had a significantly reduced MEF-senescent signature, reminiscent of the level in ESCs (**Figure 3d**). This indicates the transformed lines are avoiding senescence. Interestingly, the reprogramming-competent signature (defined in **Figure S1c**) declines in all lines (except SV40T), which suggests a reduced capability to reprogram (**Figure 3d**).

**Figure 3.**
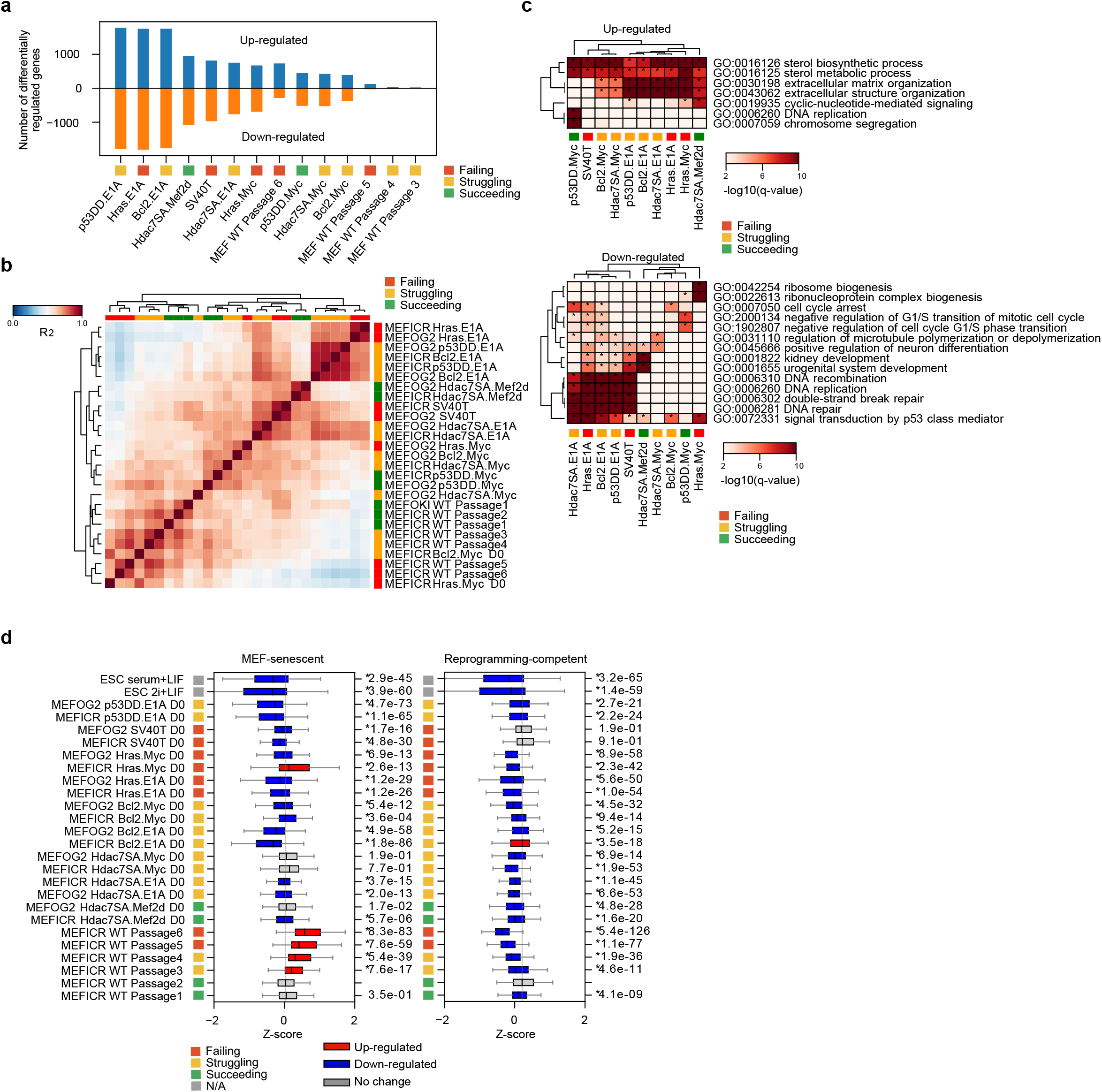
The immortalized MEFs are widely divergent from wild-type MEFs. **a** Chart showing the number of significantly differentially regulated genes up (+) or down (-) regulated (Bonferroni-Hochberg corrected p-value of <0.01 and a minimum fold-change of 2). **b** Correlation (R^2^) of the immortalized MEF lines compared to the wild-type MEFs. **c** Selected GO terms from the significantly up or down-regulated genes from the transformed MEF lines. A term was considered significant if it had a Bonferroni-Hochberg corrected p-value of <0.01). See also **Figure S4b** for the full table. **d** Boxplots for the genes defined as the MEF-senescent signature or the reprogramming-capable signature from **Figure S1c**. Significance is from a two-sided Welch’s t-test for pair-wise comparisons versus MEFICR wildtype Passage 2 as the reference. A change was considered significant if the p-value was < 0.01. Significant changes are colored red or blue.

Surprisingly, the patterns of the gene expression changes were uniform across cell lines; genes up or down-regulated in one transformed line tended to be either unchanged or similarly deregulated in other lines (**Figure S4c**). This implies a partially shared transformation signature that defines a spectrum of severity measured by the total number of differentially expressed genes.

We next looked at specific genes that have been identified as key blocks or promoters of the reprogramming process. Three factors related to cancer and cellular transformation that also modulate reprogramming in wild-type cells are *Tp53, Cdkn1a*, and *Rb1* (Retinoblastoma) [40, 50-53]. There was little change in *Rb1* mRNA levels, and whilst *Cdkn1a* was relatively high in all cell lines except p53DD.E1A, SV40T, and wildtype MEFs, its expression was not correlated with reprogramming capability (**Figure S5a**). For *Tp53*, its expression levels were relatively consistent across the MEF lines and did not correlate with the reprogramming ability of the lines (**Figure S5a**). We inferred p53 activity by looking at known target genes of p53, and all lines showed unaltered p53 activity except for the MEF line containing the dominant-negative p53DD (p53DD.E1A), which had significantly down-regulated p53 target genes (**Figure S5b, c**). Loss of p53 is beneficial for reprogramming in wildtype cells [40, 53, 54], yet in transformed lines reduced p53 activity in the p53DD.E1A line did not correlate with efficient reprogramming (**Figure 2b**). This suggests that, in contrast to wild-type cells, reduced p53 activity is not beneficial for reprogramming in transformed cells.

### Immortalized MEFs have diverse defects in reprogramming phases

Reprogramming occurs in defined phases [14, 15, 55], so we explored if there are also changes in the phases of reprogramming in the transformed lines. We performed RNA-seq during reprogramming in the early phase (D3, D6), the mid-phase (D9, D12), and the late phase (D15). These time points roughly correspond to three waves of gene expression labeled initiation, maturation, and stabilization [15, 55]. The same waves were observed in our data, based on sets of genes specific to the three phases (**Figure 4a, S6a**). Interestingly, different transformed lines completed different phases of reprogramming and appeared to fail at distinct stages. Lines that can successfully be reprogrammed: Hdac7SA.Mef2d, Bcl2.Myc and Hdac7SA.Myc, closely resembled the wild-type MEF reprogramming process (**Figure 4a**). Hdacs7SA.E1A and p53DD.E1A progressed through maturation but struggled to up-regulate stabilization phase genes (**Figure 4a**), although both can ultimately generate GFP+ reprogrammed cells at very low efficiency (**Figure 2b**). For the transformed lines that failed to reprogram, they failed at different stages. All cell lines failed stabilization, SV40T and Hras.E1A failed maturation, but completed initiation, whilst Hras.Myc failed all three stages (**Figure 4a, S6**).

The mesenchymal-epithelial transition (MET) is a key part of the early stage of reprogramming [55, 56]. We checked that the transformed MEFs remained mesenchymal-like, and they still expressed typical markers of the mesenchyme (**Figure 4b**). All transformed lines managed to navigate the MET, except for Hras.Myc (**Figure S6b**). Indeed, some lines, including two transformed lines that would ultimately fail to reprogram; Hras.E1A and SV40T, up-regulated epithelial genes in advance of wild-type reprogramming cells (**Figure S6b**), including two transformed lines that would ultimately fail to reprogram, Hras.E1A and SV40T. This suggests that failure to execute a MET affected only the Hras.Myc line.

**Figure 4.**
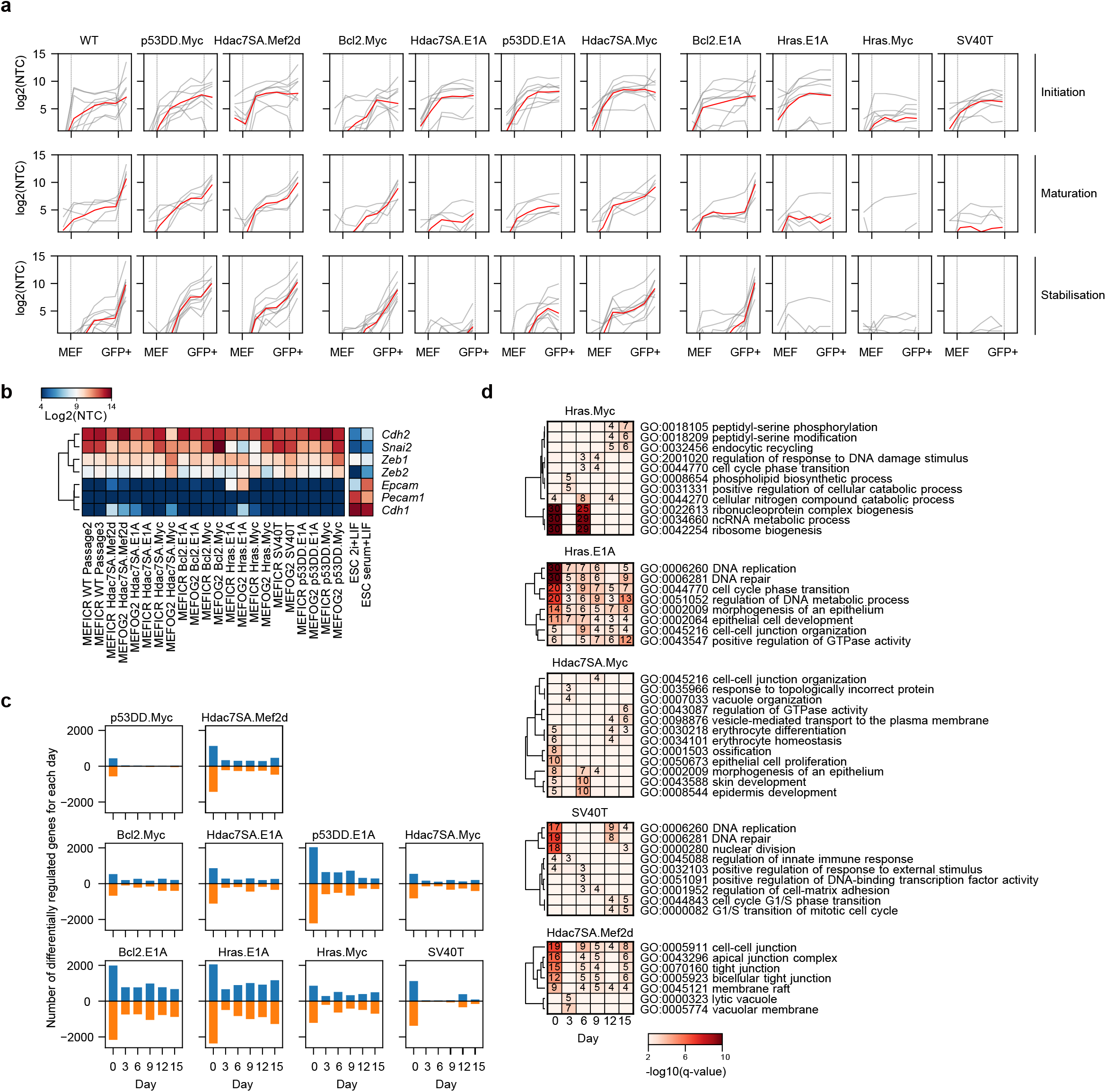
Transformed MEFs fail at different phases of reprogramming. **a** Line charts showing the expression of initiation, maturation, and stabilization genes [55], at the indicated time points or in MEFs and GFP+ cells. See **Figure S6** for the expression of the individual genes in each class. The red line is the mean of all genes, the grey lines are the individual genes. **b** Heatmap of the expression of mesenchymal and epithelial marker genes in starting lines and ESCs. **c** Bar charts showing the number of significantly differentially regulated genes at each time point for the indicated transformed lines versus a wildtype reprogramming time course. Differential expression was compared for each line and time point versus the corresponding wildtype timepoint. A gene was considered differentially regulated if the q-value (Bonferroni-Hochberg corrected p-value) was <0.01 and the fold-change >2. **d** Heatmaps of significantly overrepresented gene ontology (biological process) terms for down-regulated genes from the indicated time points and cell lines.

We next looked at changes in the overall phases of reprogramming in the early, middle, and late stages of reprogramming. We measured the number of significantly deregulated genes at each time point and used this as a proxy for the divergence from a typical reprogramming time course. In this analysis, down-regulated genes represent genes that fail to be correctly up-regulated at the specific day of the time course, whilst up-regulated genes those represent those genes that are erroneously high during reprogramming (**Figure 4c**). A large number of differentially regulated genes on a specific day represents a divergence from the expected reprogramming stage. As expected, the lines that reprogrammed at the highest efficiency (p53DD.Myc) also had the lowest overall divergence (**Figure 4c**). Lines that could reprogram, but at low efficiency (Hdac7SA.Mef2d, Bcl2.Myc, Hdac7SA.E1A, and Hdac7SA.Myc) tended to have high initial divergence from the MEF state (day 0), but would correctly regulate reprogramming-associated genes at the later time points. Finally, lines that fail to reprogram (Hras.E1A and Hras.Myc), or reprogram exceptionally rarely (Bcl2.E1A), tended to have high divergence at all time points. The exception was the SV40T line, which appeared to follow the reprogramming gene expression program closely, but would diverge at day 12. Interestingly, GO analysis of the down-regulated genes (genes that should be up-regulated at that specific timepoint) highlighted the regulation of DNA repair genes, which was defective in both Hras.E1A and SV40T cell lines (**Figure 4d**). However, overall, there was a surprising diversity of changes in the transformed MEF lines that give rise to a range of defects at specific time points of the reprogramming process. This data indicates that transformation-specific effects drive line-specific blocks on reprogramming in a stage-dependent manner.

### Chemical intervention can convert some lines from reprogramming-incapable to reprogramming-capable

The analysis above suggested several pathways that could be manipulated, specifically, the MET, ribosome biogenesis, cell proliferation, cell adhesion, DNA repair, and apoptosis. We wondered if it was possible to intervene in the reprogramming process and convert the reprogramming-incapable to -capable. To investigate this we assayed the effect on reprogramming of a range of inhibitors on two reprogramming-capable lines (WT and Hdac7SA.Mef2d MEFs) compared to two reprogramming-incapable lines (Hras.E1A and SV40T MEFs). In total, we used 25 inhibitors targeting a range of pathways (**Figure S7a**). Whilst most inhibitors had no effects or would ablate reprogramming, we observed that 5 inhibitors promoted the reprogramming of one or more of the transformed reprogramming-incapable lines (**Figure S7a**). The five small molecules target: MEK1/2 (PD; PD0325901), GSK3 (CHIR; CHIR99021), ROCK (Y; Y23637), G9a (BIX; BIX-01294), and histone deacetylases (TSA). We tested the inhibitors in combinations and found that the most efficient combination was CHIR, PD and Y (**Figure 5a, S7b**). However, only PD was essential to convert SV40T and Hras.E1A to reprogramming-capable, as serum+Vc+PD alone resulted in a small number of GFP+ cells. Conversely, CHIR, Y, and BIX alone were incapable of converting Hras.E1A MEFs to reprogramming-capable (**Figure S7b**). Interestingly the inhibitor cocktails had effects on reprogramming in a transformed line-specific manner, as the addition of PD to the reprogramming cocktail converted the normally reprogramming-capable Bcl2.E1A line to incapable. This highlights the line-specific nature of the reprogramming barriers and shows that overcoming a barrier in one line can potentially initiate a barrier in another transformed line.

**Figure 5.**
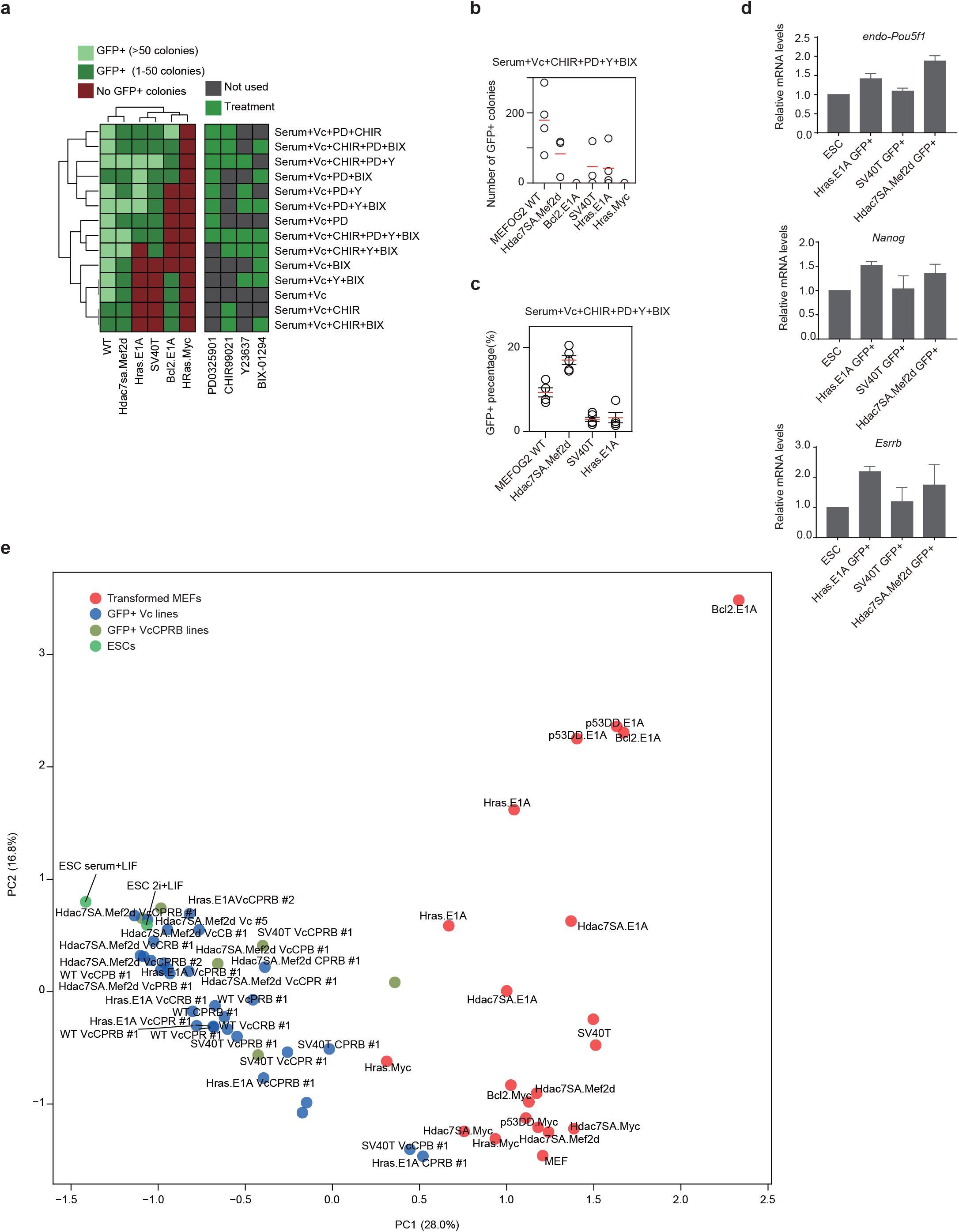
Chemical inhibition can convert some reprogramming-incapable immortalized MEFs to reprogramming-capable. **a** Heatmap showing the number of GFP colonies produced depending upon the reprogramming conditions used, for the indicated transformed lines. **b** Chart showing the number of GFP+ colonies in the indicated immortal cell lines with the inhibitor cocktail. Data is from four biological replicates (MEFOGF WT) or from three biological replicates (all other transformed cell lines). **c** Percentage of cells sorted by FACS at day 12 of the reprogramming assay in the indicated transformed cell lines or wildtype MEFs. Data is from four biological replicates. **d** RT-qPCR at day 15 of a reprogramming experiment using primers targeting endogenous Oct4 (endo-*Pou5f1*), *Nanog, and Esrrb*. The mean of three biological replicates ± s.d. is shown. **e** PCA of the RNA-seq from the immortalized MEFs, and the reprogrammed iPSCs and ESCs. The various chemical cocktails used during the reprogramming are abbreviated for clarity. Key: Vc=Vitamin C; P=PD0325901; C=CHIR99021; R=Y23637; B=BIX-01294.

We confirmed the resulting iPSC-like cells derived from Hras.E1A or SV40T MEFs reprogrammed with the PD, CHIR, Y and BIX adopted a normal morphology, expressed pluripotent marker genes by qRT-PCR and Western blot (**Figure 5d, and Figure S7c**. We sorted the GFP+ cells and established iPSC lines, and measured their gene expression which closely correlated with ESCs by correlation and PCA (**Figure 5e and Figure S8a, b**). It should be emphasized that the chemical cocktail did not improve (or inhibit) wildtype MEF reprogramming, as wildtype cells reprogrammed with similar rates in serum+Vc or the inhibitor cocktail (**Figure S7b**). This emphasizes that the blocks seen in transformed cells are specific to transformation and are not present in wild-type untransformed cells. Overall, the results indicate that transformation-specific pathways are impairing the ability to reprogram.

### Epigenetic defects underly the inability to reprogram

We next focused on three specific cell lines: Hdac7SA.Mef2d, Hras.Myc and Hras.E1A. Hdac7SA.Mef2d could reprogram in serum under normal conditions, whilst Hras.E1A is initially resistant to reprogramming (**Figure 2b**), but the addition of PD converts it to reprogramming-capable (**Figure 5a-d**). Hras.Myc, conversely, could not be converted to reprogramming-capable with any of the conditions we tried (**Figure 5b**).

We explored the epigenetic regulation of the transformed MEFs. As a proxy for overall epigenetic activity, we looked at the expression levels of epigenetic factors involved in activation, repression, and the reading of epigenetic marks, as defined by the Epifactors database [57]. As expected, epigenetic-related factors were uniformly significantly up-regulated in ESCs compared to MEFs (**Figure S9a**), reflecting their more complex epigenetic regulation [58]. For the transformed MEFs, activators were more often significantly up-regulated, compared to untransformed MEFs (**Figure S9a**). Erasers and readers were only up-regulated in several transformed lines, all containing E1A: Hras.E1A and Bcl2.E1A and p53DD.E1A (**Figure S9a**). Nonetheless, whilst the different classes of epigenetic regulators varied across the transformed lines and did not discriminate reprogramming-capable from incapable, the general pattern for the majority of lines was increased expression of epigenetic regulators. Western blot of repressive histone modifications (e.g. H3K27me3, H3K9me3) tended to stay the same in the different lines (**Figure 6a**). However, histone acetylation inversely correlated with reprogramming capability as H3K27ac, H3ac or H4ac was high in reprogramming-incapable lines (Hras.Myc, Hras.E1A and SV40T). The results support epigenetic deregulation as a feature of the transformed lines and suggest increased epigenetic activation through histone acetylation is a feature of transformed reprogramming-incapable MEFs.

**Figure 6.**
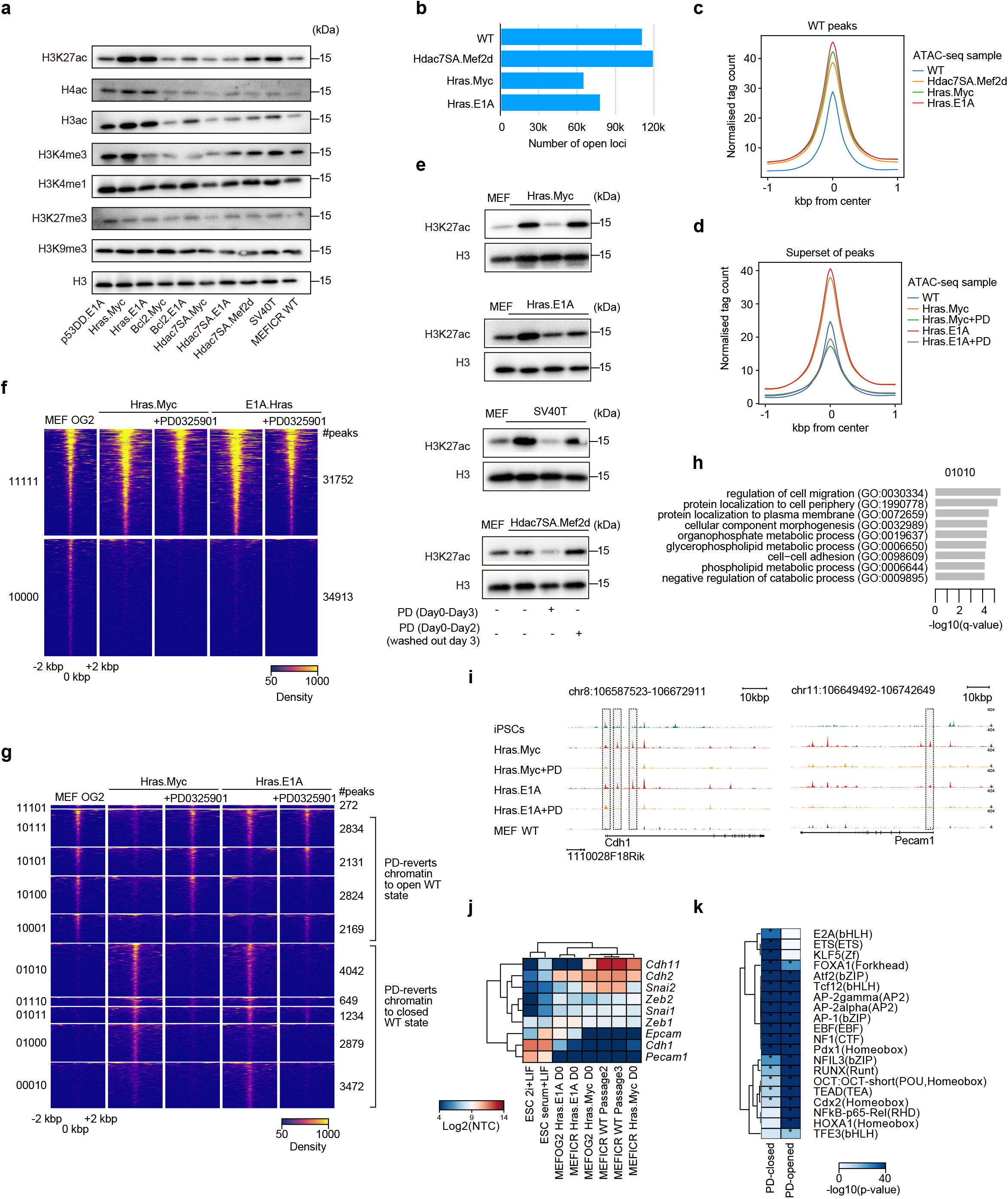
Inhibition of MEK repairs some epigenetic defects in the transformed MEFs. **a** Western blot for a selection of epigenetic marks and total histone H3 as a loading control. **b** Bar chart showing the number of ATAC-seq loci identified as open/accessible in the indicated wildtype or transformed MEFs. **c** Pileups of the ATAC-seq data centered on all wildtype MEF peaks and showing the flanking 1 kbp (kilo base pairs). Normalized tag counts (reads per million per bin) are shown for the indicated samples. **d** Pileups of the ATAC-seq data centered on a non-redundant superset of all peaks from all of the ATAC-seq samples. The plot is centered on the peak summit and shows the flanking 1 kbp. Normalized tag counts are shown for the indicated samples. **e** Western blot of activatory histone marks upon addition of PD to the MEFs for 3 days, or cells treated with PD for two days and then washed out on the third data (washed out on day 3). **f** Heatmap of the ATAC-seq data showing the permanently open (11111), and open only in the wildtype MEFs (10000). The number of peaks in each class is indicated on the right-hand side. Read density is shown centered on the peak summit and 2 kbp on each side. The full heatmap containing all groups is in **Figure S9b**. **g** Heatmap of selected peak groups for the indicated ATAC-seq samples. The full heatmap containing all groups is in **Figure S9b**. **h** Gene ontology analysis for nearby genes (a TSS within 10 kbp of an open locus) from the 01010 group (closed in wildtype MEFs, PD causes these loci to close in the transformed MEFs) from **Figure 6g**. **i** Genome pileup view of the ATAC-seq data showing the *Cdh1* and *Epcam* loci in wildtype MEFs, Hras.E1A, and Hras.Myc MEFs with or without treatment with PD. Genome assembly is mm10. Regions of open chromatin that are modified in the PD-treated cells are indicated by vertical grey bars. **j** Expression heatmap of selected epithelial and mesenchymal genes in the indicated MEF lines and ESCs. **k** Motif analysis of selected PD-close and PD-open genes.

### Transformed MEFs have a hyperactive chromatin state which MEK-inhibition partially reverts to wildtype

To explore changes in the epigenetic state of the MEFs we performed ATAC-seq on three transformed lines: Hdac7SA.Mef2d (reprogramming-capable), Hras.E1A (initially incapable, but can be induced to reprogram with MEK inhibition), and Hras.Myc (reprograming-incapable) and compared them to wildtype MEFs. Heatmaps of open and closed chromatin indicated, surprisingly, that the Hdac7SA.Mef2d lines were distal from all of the other transformed cell lines and had the greatest number of specific peaks (**Figure S9b, Figure 6b**). The loci that were opened or closed in the Hdac7SA.Mef2d lines were significantly correlated with matching changes in gene expression (**Figure S9c**). Intriguingly, the overall ATAC signal was substantially higher in all three transformed cell lines compared to wildtype MEFs (**Figure 6c**), and this effect was despite fewer open loci in the Hras.Myc and Hras.E1A lines (**Figure 6b**). This suggests that the transformed MEFs have increased levels of open chromatin.

We next focused on Hras.Myc and Hras.E1A lines, and removed the Hdac7SA.Mef2d from further analysis as it is capable of reprogramming under normal conditions and its ATAC-seq genome-wide pattern was distal from the other transformed cell lines and wildtype MEFs. Hras.Myc and Hras.E1A are interesting as they have relatively similar ATAC-seq patterns (**Figure S9b**), yet Hras.E1A can be converted to reprogramming-capable, whilst Hras.Myc cannot. The two lines have Hras, which does not impede reprogramming independently. Similarly, other lines co-transformed with Myc are also reprogramming-capable (Bcl2.Myc and Hdac7SA.Myc), as E1A transformed lines (Hdac7SA.E1A). Hence, the individual transgenes are not incompatible with reprogramming, only the specific combinations of Hras.E1A and Hras.Myc is important. One key difference between Hras.E1A and Hras.Myc is that only Hras.E1A up-regulated initiation-stage genes and underwent an MET (**Figure 4a, d**), suggesting the impairment of reprogramming is already primed in the starting transformed MEFs.

To explore the underlying differences, we performed ATAC-seq on Hras.Myc and Hras.E1A lines with and without PD treatment, and compared them to wildtype MEFs. Interestingly, the global upregulation of the ATAC-seq signal at open loci was lost in the PD-treated samples and returned to levels similar to the wild-type (**Figure 6d**). Western blot of H3K27ac, a mark of open active chromatin, supported this observation as H3K27ac was higher in the transformed MEFs, and treatment with PD for 3 days reduced H3K27ac to wildtype levels (**Figure 6e**). Interestingly the result is transient, as treatment for 2 days followed by washing out reverted H3K27ac levels to the hyperactive state (**Figure 6e**). This suggests that when PD inhibits MEK it leads to the indirect closing of chromatin and reductions in the global levels of H3K27ac. However, this effect was transient, and incomplete as the overall epigenetic landscape did not revert to a wildtype state and 34,913 loci open in wildtype MEFs remained closed in the transformed cell lines (**Figure 6f**).

To explore the mechanisms underlying the action of PD, we next divided the loci into two types based upon their state in wildtype MEFs and the effect of PD: (1) those loci that PD opens and reverts to a wildtype state, and (2) those loci that PD closes, reverting to the wildtype state (**Figure 6g**). There was a roughly equal split between the two classes (10,230 peaks versus 12,276 peaks; **Figure 6g**). GO analysis of the peaks that were specifically reverted to wildtype state in the Hras.Myc cells suggested that cell-cell adhesion gene loci were being remodeled (**Figure 6h**). Indeed, chromatin at *Cdh1* and *Epcam* (two epithelial genes) were open in the Hras.E1A and Hras.Myc MEFs, but were closed when the cells were treated with PD and reverted to a wild-type MEF pattern at these genes (**Figure 6i**). Importantly, *Cdh1, Epcam*, and other epithelial genes are not expressed in the transformed MEFs (**Figure 6j**), indicating that epigenetic regulation at epithelial gene loci is deleterious for reprogramming of transformed cell lines.

We next looked at possible downstream targets of MEK signaling that mediated this indirect effect on chromatin. We used motif analysis to look for changes in transcription factor occupancy at PD-modulated peaks (**Figure 6k**). Interestingly, peaks that were closed in response to PD treatment were specifically enriched with ETS, E2A, and KLF motifs. KLF proteins are major regulators of MET processes and are pluripotent factors, whilst ETS factors have been identified as barriers for wild-type untransformed reprogramming [59]. Loci that PD treatment opens were enriched with REL, HOX, and TFE family TFs (**Figure 6k**). To explore deeper we looked at two small groups of chromatin loci that distinguished reprogramming-capable from -incapable, 10001 (specifically open in WT and Hras.E1A+PD) and 01110 (specifically closed in WT and Hras.E1A+PD) (**Figure 6g**). Interestingly, GO analysis of the 01110-group suggested that enhancers near to genes related to cell differentiation were erroneously opened (**Figure S10a**), for example at the developmental-related genes *Isl1* and *Tcf7l2* (**Figure S10b**). This was a purely an effect on chromatin state as *Isl1* and *Tcf7l2* were below the threshold for detection in the RNA-seq data. This pattern extended to all of the genes in the vicinity of the 01110-chromatin state which were not significantly up or down-regulated (**Figure S10c**). Conversely the genes associated with the 10001 group were significantly down-regulated (**Figure S10c**), although GO analysis suggests they are related to basic cellular processes such as cytoskeleton organization (**Figure S10d**). Overall, this suggests the epigenetic activation of developmental genes is deleterious for reprogramming, and MEK inhibition can indirectly modulate the epigenetic state to close chromatin.

## Discussion

Reprogramming cancerous/transformed cells to pluripotent cells, followed by their differentiation, could form a valuable experimental model to recapitulate the earliest stages of tumorigenesis [60, 61]. However, reprogramming of primary cancerous tissues is often difficult or impossible for many cancers, and even in possible cases the efficiency is considerably less than wild-type cells. Additionally, there remains doubts about the *bona fide* reprogramming of the cancerous cells [28, 34, 35], and it is not clear if reprogramming of primary human cells is complete. A significant problem is the heterogeneity of primary cancer cells. To overcome this problem, we focused on isogenic transformed MEF lines that allow us to control overall heterogeneity. Although, even in this system there was a surprisingly large variation between the different methods of MEF transformation.

There is an intimate but unclear link between reprogramming and transformation [1, 61]. This link was recently explored in a novel system with multiple inducible genes that could be manipulated to either transform MEFs or perform reprogramming to iPSCs [17]. This elegant study showed that reprogramming and transformation followed similar initial paths, but then later diverged. The system they used for transformation was K-ras^G12D^ and *Myc* overexpression [17], which is similar to the Hras.Myc (*Hras*^*G12V*^/*Myc*) line described in this manuscript. Intriguingly, we show that Hras.Myc is not compatible with reprogramming, and those cells fail very early in the reprogramming process. However, several other lines described here can be transformed and remain compatible with reprogramming. Potentially, transformation strategies that remain compatible with reprogramming may recapitulate more of the reprogramming process as they transform, and only diverge from reprogramming at later stages.

*Ras* overexpression is beneficial for reprogramming wildtype MEFs, but is a major barrier to reprogramming transformed cells [62]. Indeed, when Hras^G12V^ was overexpressed in p53-null immortalized MEFs reprogramming was blocked [62]. This is reminiscent of the effect we observed in the Hras.Myc and Hras.E1A lines, which are also resistant to reprogramming. However, we show that inhibition of MEK by PD can only enable Hras.E1A cells to reprogram, and Hras.Myc cells remain reprogramming-incapable. This indicates that Hras^G12V^ is context-dependent when its co-transformation factor is different.

Several studies have identified small molecules that assist in reprogramming transformed cells to a more pluripotent-like state. mTOR was also identified as a barrier to the reprogramming of sarcoma cells [63]. In our experiments, mTOR inhibition had no effect, and ROCK inhibition worked only to boost reprogramming in cooperation with MEK inhibition. This again highlights the context-specific nature of the barriers in different transformed cell lines. Overall, MEK inhibition was the dominant factor, which agrees with a previous study that showed that MEK inhibition assisted in reprogramming human transformed cell lines [64]. PD is also a key component of the 2iLIF medium used to maintain mouse ESCs in a naïve groundstate [49, 65]. It is unclear if PD plays the same role in assisting in reprogramming transformed MEFs and in promoting pluripotency. Indeed, reprogramming with PD is deleterious for reprogramming wild-type MEFs, at least in the early stages of reprogramming [62, 66].

Ultimately, these studies support a model for the context-specific reprogramming of transformed cell lines. Different transformations lead to diverse blocks at different stages of the reprogramming process. Inhibition of ERK by PD can overcome some of these barriers, primarily in Ras-transformed cells [62, 64]. However, each transformed cell line can harbor defects that block reprogramming at unique stages. This perhaps reflects the diversity of regulatory features disrupted in cancer and may relate to the ‘stemness’ of a cancer type or method of transformation [67]. Whilst there are many surprising similarities between tumorigenesis and reprogramming, the nature of the relationship remains unclear [1]. Potentially, insights into reprogramming can inform the process of transformation and provide new models to address tumorigenesis and a clearer understanding of cell type plasticity in cancer development.

## METHODS SUMMARY

### Animal experimental approval

Ethical approval for the generation of MEFs and teratoma assays was granted by the Southern University of Science and Technology animal ethical committee, approval number: SUSTC-2019-005.

### Cell culture and MEF transformation

MEFs were derived from E13.5 embryos from OG2 mice [36], or a wild-type ICR genetic background. MEF cells were cultured in DMEM high-glucose media containing 10% FBS (GIBCO), 1 × GlutaMAX (GIBCO), and 1x nonessential amino acids (GIBCO), 0.5x Penicillin/Streptomycin (Hyclone). We generated immortalized MEF cell lines by transfecting the following transgenes singly or in combination: P53DD, PMM2-SV40T, PMM2-Hdac7SA, PMM2-Mef2d, Bcl2, PMM2-Myc, Ras, PMM2-E1A.

Immortalized MEFs were passaged for at least a month to remove wild-type untransformed MEFs.

### Somatic cell reprogramming

Wildtype MEFs and transformed MEFs in an OG2 reporter background were reprogrammed as described in [36]. Briefly, OKSM in lentivirus was transfected into 15,000 MEFs in one well of a 12-well plate. One day after transfection the medium was changed to reprogramming medium: DMEM high-glucose media containing 15% FBS (GIBCO), 1× GlutaMAX (GIBCO), 1x non-essential amino acids (GIBCO), 0.5x Penicillin/Streptomycin (Hyclone), 1mM sodium pyruvate (GIBCO), 0.1 mM 2-mercaptoethanol (GIBCO), 1000 U/ml leukemia inhibitory factor (LIF) (Millipore) and vitamin C (50 μg/ml, Sigma), as indicated in the figures. The medium was changed daily. Inhibitors were added when the medium was changed to reprogramming medium: PD0325901 (1 μM, Sigma), CHIR99021 (3 μM, Sigma), Y23637 (10 μM, Sigma), BIX-01294 (200 nM, Selleck), and TSA (50 nM, Sigma).

### Karyotype analysis

When the immortalized MEF cell growth density reached 80 to 90%, colchicine (0.2 μg/ml, Selleck), was added to the culture medium to a final concentration of 0.2 μg/ml. The cells were incubated at 37°C for 120 min. After colchicine treatment, cells were washed twice with PBS, and 0.5 ml of 0.25% trypsin was added for digestion. Then, 7 ml of 0.075 mol/L KCl solution preheated to 37°C was added, and the cell suspension was blown with a straw and incubated at 37°C in a water bath for 25 mins. Collected cells were fixed with a fixing solution (methanol glacial acetic acid 3:1 preparation) at 37°C, 3 min. After fixation, cells were centrifuged at 1200 revolutions per min (rpm) for 5 min, after which the supernatant was discarded. 7 ml of fresh fixing solution was added, and the cells were gently beaten with a pipette tip, and fixed in a 37°C water bath for 40 mins. Cells were centrifuged at 1200 rpm for 5 min, after which the supernatant was discarded. Subsequently, 7 ml of fresh fixing solution was added, followed by gentle beating with a filter tip. The cells were fixed in a 37°C water bath for 40 min. After fixation the cells were centrifuged at 1200 rpm for 5 min, and then most of the fixation solution was removed, and the cells were resuspended with part of the fixation solution. Suspended cells were pipetted onto a slide from a distance of about 30 cm. Immediately after dropping the cells on the slide, move the slide into the oven at 75°C and bake for 3h. Next, 0.03 g trypsin powder to 55 ml normal saline was added, followed by a gentle shaking, before adjusting the pH to about 7.2 with 3% Tris-HCl. The preparation was put into trypsin digestion solution for 8-10 seconds, followed by the addition of saline to stop digestion, then add Giemsa staining (C0133, Beyotime Biotech) solution was added for 5-10 minutes, and then pick out the glass slide with tweezers. Both sides were gently rinsed with running water and allowed to dry at room temperature. After the slides dried, they were examined under a microscope to look for a good chromosome split.

### Teratoma formation

5-10 million mouse PSCs in a slurry of Matrigel and mTeSR (1:1) medium were injected into 8-week-old female nude (BALB/cNj-Foxn1^nu^/Gpt) immunodeficient mice. Teratoma growth was quantified by measuring approximate elliptical area (mm^2^) with calipers measuring the outward width and height after growth for 60 days. Representative tumors were dissected and sectioned and slices were stained with hematoxylin and eosin.

### Alkaline phosphatase staining

Reprogrammed cells were stained with alkaline phosphatase according to the manufacturer’s protocols. In brief, reprogramming cells were fixed in 1% (w/v) formaldehyde, and then cells were stained with BCIP/NBT Alkaline Phosphatase Color Development Kit (C3206, Beyotime Biotech) according to the kit’s instructions.

### Flow cytometry

Reprogramming cells were digested with trypsin and washed with DPBS once, and analyzed or sorted with BD FACS Aria III flow cytometer. To monitor cellular reprogramming status, reprogrammed cells were stained with antibodies: anti-CD44-PE (25-0441-82, 1:500, Thermo Fisher) and anti-ICAM-1-FITC (sc-8439 FITC, 1:200, Santa Cruz Biotechnology). Cells were sorted and analyzed on a BD FACS Aria III instrument.

### Western blot

Cells were lysed with RIPA buffer (P0013B, Beyotime Biotech). Proteins were resolved in sodium dodecyl sulfate-polyacrylamide gel electrophoresis (SDS-PAGE) and transferred onto pre-activated polyvinylidene fluoride (PVDF) membranes (IPVH00010, Millipore, MA, USA). The PVDF Membranes were incubated with anti-H3 (ab1791, 1:3000, Abcam), anti-H3K27ac (ab4279, 1:2000, Abcam), anti-H3ac (06-599,1:2000, Millipore), anti-H4ac (ab46983, 1:2000, Abcam), anti-H3K4me3 (ab8580, 1:2000, Abcam), anti-H3K4me (ab176877,1:2000, Abcam), anti-H3K27me3 (07-449,1:2000, Millipore), anti-H3K9me3 (ab8898, 1:2000, Abcam), anti-NANOG (8822S, 1:1000, cell signaling), anti-OCT4 (sc-5279, 1:1000, Santa Cruz Biotechnology), anti-P53 (sc-126, 1:1000, Santa Cruz Biotechnology), anti-HDAC7 (ab12174,1:2000, Abcam), anti-MEF2D (ab32845,1:2000, Abcam), anti-ACTIN (ab8227, 1:3000, Abcam), anti-MYC (ab32074,1:2000, Abcam), anti-SOX2 (4900S, 1:1000, cell signaling). Afterward, the membranes were incubated with HRP-conjugated goat antirabbit IgG (ab205718, 1:3000, Abcam) and visualized using an enhanced chemiluminescence BeyoECL method (P0018AS, Beyotime Biotech).

### Immunofluorescence

Cells were washed with cold PBS and fixed using 4% (w/v) paraformaldehyde in PBS for 10 minutes at room temperature. Then cells were washed with PBS and permeabilized with 0.25% (v/v) Triton X-100 for 10 minutes at room temperature. Fixed cells were then washed with PBS three times and blocked with 3% (w/v) bovine serum albumin for 1 hour at room temperature. Then cells were incubated with primary antibodies: anti-NANOG (8822S, 1:1000, cell signaling), at 4°C overnight in primary antibody. Following overnight incubation, cells were washed with wash buffer (0.1% (v/v) Tween-20 in PBS) and incubated with secondary antibodies Alexa Fluor series (Life Technologies) in wash buffer for 1 hour at room temperature. Cells were washed again and nuclei were stained using DAPI (5 μg/ml, Thermo).

### RT-qPCR

Total RNA from cells was isolated using RNAzol RT (MRC, RN190) according to the manufacturer’s protocols. cDNA synesis by using a PrimeScript RT Master Mix (Takara, RR036A). Real-time PCR was performed in triplicate using SYBR Premix Ex Taq (Takara, RR820A) and using a Biorad Real-time PCR system. The primers used are listed in **Supplementary Table S2**.

### RNA-seq preparation and analysis

RNA was isolated using RNAzol RT (MRC, RN190) according to the manufacturer’s protocol and prepared for sequencing with RNA-seq NEB Next Ultra RNA Library Prep Kit (NEB, #7530). Samples were sequenced on an Illumina Novaseq 6000.

RNA-seq was analyzed essentially as described in [47], except scTE/te_counts was used to assign mapped reads to genes and TEs [68], and the UCSC genome browser repeatmask track and GENCODE vM23 annotation were used for TE and gene assignment. Differential expression was determined using DESeq2 [69]. A gene was considered differentially expressed had an absolute fold-change of at least 2, and a Bonferroni-Hochberg corrected p-value of 0.01. GSEA was performed using fgsea [70], and GO was done with goseq [71]. Cell type determination was scored with DPre [48]. Other analyses were performed using glbase3 [72].

### ATAC-seq preparation and analysis

ATAC-seq library was generated using the Tn5 enzyme from the TruePrep DNA Library Prep Kit V2 (Vazyme, TD501-02) as previously described [73]. Briefly, a total of ∼50,000 cells were washed once with 50 μl of cold PBS and resuspended in 50 μl lysis buffer (10 mM Tris-HCl pH 7.4, 10 mM NaCl, 3 mM MgCl2, 0.2% (v/v) IGEPAL CA-630). The nuclei were centrifuged for 10 min at 500 × g at 4 °C, followed by the addition of 50 μl transposition reaction mix (25 μl TD buffer, 2.5 μl Tn5 transposase (Vazyme), and 22.5 μl nuclease-free water). Samples were PCR-amplified and purified using a MinElute kit (Qiagen). After selecting an appropriate PCR cycle number (See [73]) samples were sequenced on an Illumina sequencer. ATAC-seq data was analyzed essentially as described in reference: [59]. Briefly, reads were aligned to the mm10 mouse genome with bowtie2 [74], peaks were called with MACS2 [75], and then the *redefine_peaks* function, which is a generalized reimplementation of the algorithm in [59], was used to recover low-scoring peaks by sharing peak information across samples [76]. DNA binding motifs were detected using HOMER [77]. All other analyses were performed using glbase3 [72]. ATAC-seq data from GSE93029 [59] and GSE103980 [78] were reanalyzed as part of this study.

### Statistical analysis and biological replication

Differential gene expression was calculated using DESeq2 (v1.36.0). A gene was considered significantly differentially regulated if it had an absolute fold-change of at least 2 and a Bonferroni-Hochberg corrected p-value (q-value) of <0.01. This criterion was used in **Figures 3a, 4c, Figures S1c, and d**. RNA-seq experiments were performed in at least biological duplicate (different samples on different days, or independent cell lines). Gene ontology analysis was performed using goseq (v1.48.0) and statistics were calculated using goseq’s internal statistical model. A gene ontology category was considered significantly enriched if there were at least 50 genes in that GO term and a Bonferroni-Hochberg corrected p-value (q-value) of <0.01. GSEA was performed using fgsea (v1.22.0). Gene sets were considered enriched or depleted if they had an absolute NES (normalized enrichment score) of at least 1.5 and a Bonferroni-Hochberg corrected p-value (q-value) of <0.01. Western blots were repeated at least twice with similar results, with the exception of **Figure S3b** and **S7c** which were performed once. **Figure 6a** was repeated three times. FACS analysis (**Figure 1c**) was performed twice. qRT-PCR was performed on at least three biological replicates with three technical replicates each. Transcription factor motif analysis was performed using HOMER. A motif was considered significantly enriched if the uncorrected p-value was <0.00001.

## Acknowledgments

This work was supported by the National Natural Science Foundation of China (32150710521), and the Shenzhen Innovation Committee of Science and Technology (ZDSYS20200811144002008). Additional supported was rendered by the Center for Computational Science and Engineering of Southern University of Science and Technology.

## Author contributions

X.F. designed and performed the majority of the experiments, prepared the manuscript figures and revised the text. Q.Z. designed the project and planned the experiments. G.M., H.H., Y.L., J.C., Z.X., B.D., S.L. performed some experiments or assisted in experimental procedures. I.A.B., L.S., assisted with the bioinformatic analysis. R.J. assisted in manuscript writing, acquired funding and supervised H.H.. A.P.H. performed some of the bioinformatic analysis, drafted the manuscript. All authors read and revised the manuscript.

## Author information

RNA-seq data are available in the Gene expression omnibus (GEO) public database under accession number GSE213225.

