## Supplementary Material for "Modulating the epigenetic state promotes the reprogramming of transformed cells to pluripotency in a line-specific manner"

#### Supplementary Tables

**Supplementary Table 1** – Genes assigned to the 'MEF senescent signature' and the 'reprogramming permissive signature'.

**Supplementary Table 2** – Sequences of the transgenes used for aligning the RNA-seq data.

#### Supplementary Results

##### Measurement of the transgenes in the RNA-seq data

Transgene expression was measured by aligning the RNA-seq to an index composed of the FASTA sequences for the transgenes used in this study (**Supplementary Table 3**). The HsMYC (human c-MYC; ENST00000259523) measures the expression of the polycistronic OSKM transcript (Warlich et al., 2011). Note that the HsOCT4, HsSOX2 and HsKLF4 transgenes are not detectable in this RNA-seq data as the RNA is cleaved and only the HsMYC has a poly-A tail. GFP/eGFP measures the expression of the GFP reporter from the OG2 reporter (Zhuang et al., 2018).

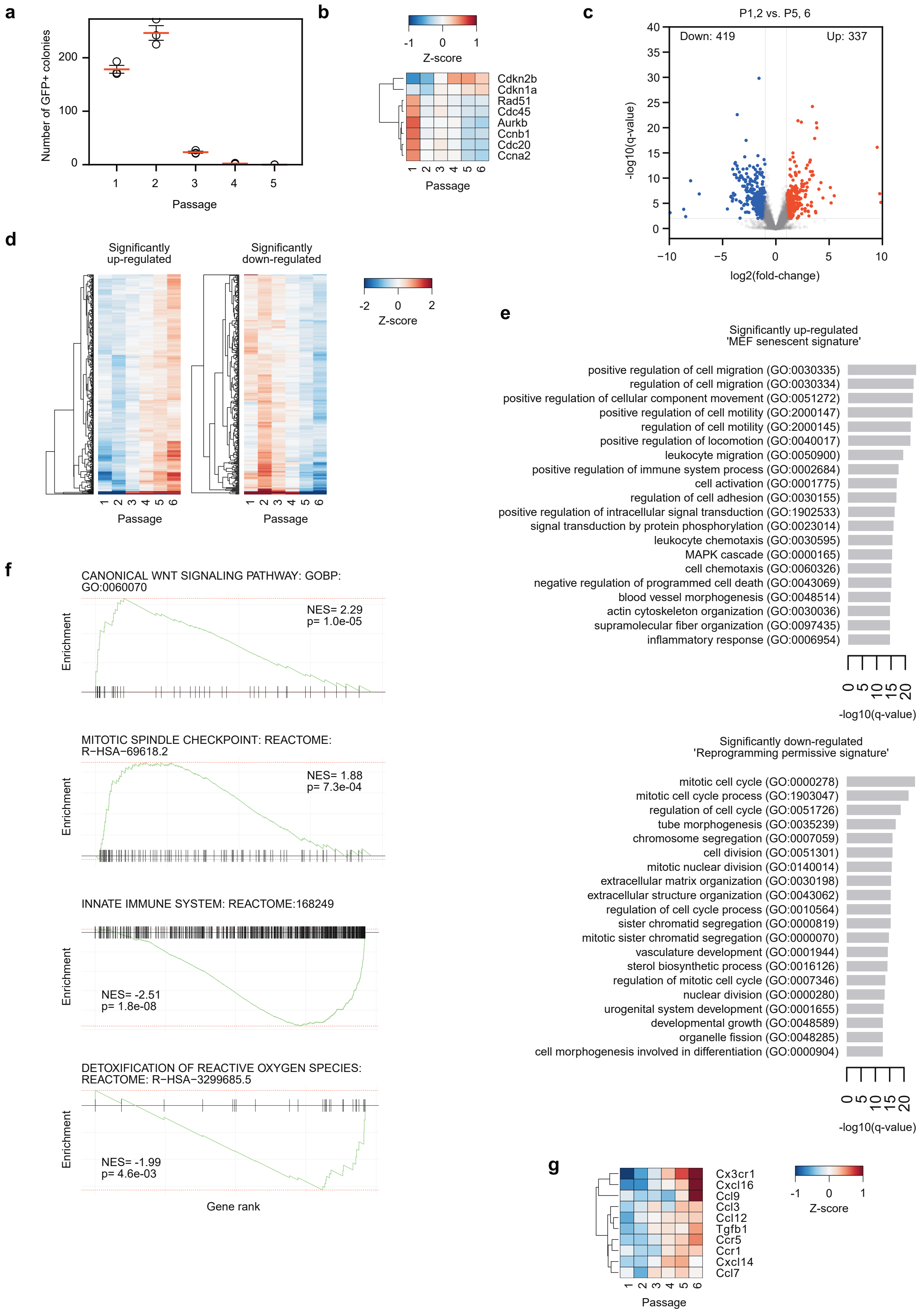


**Supplementary Figure1**

**Supplementary Fig. 1. Late passage MEFs lose reprogramming capability and become senescent.**

- **a** Number of GFP+ colonies from a reprogramming experiment using OCT4KI-MEFs from the indicated passage number. Data is from three biological replicates
- **b** Heatmap for the expression of selected cell cycle genes in the different passages of MEFs.
- **c** Volcano plot for differentially expressed genes when comparing passage 1, 2 versus passage 5, 6. A gene was considered significantly different if it had an absolute fold-change of 2 and a q-value of less than 0.01 (Bonferroni-Hochberg corrected p-value).
- **d** Heatmaps of the expression of the significantly up or down-regulated genes in the indicated MEF passages.
- **e** Gene ontology analysis of the significantly up and down-regulated genes. A GO term was considered significant at a Bonferroni-Hochberg corrected p-value of less than 0.01. Based on these GO, we labelled the up-regulated genes as the ‘MEF senescent signature’ and the down-regulated genes as the ‘reprogramming permissive signature’
- **f** GSEA for the up and down-regulated genes. Selected Reactome or GO BP (biological process) terms are shown. NES (normalized enrichment score), and the Bonferroni-Hochberg adjusted p-value are indicated.
- **g** Heatmap of selected significantly differentially expressed cytokines and chemokines.

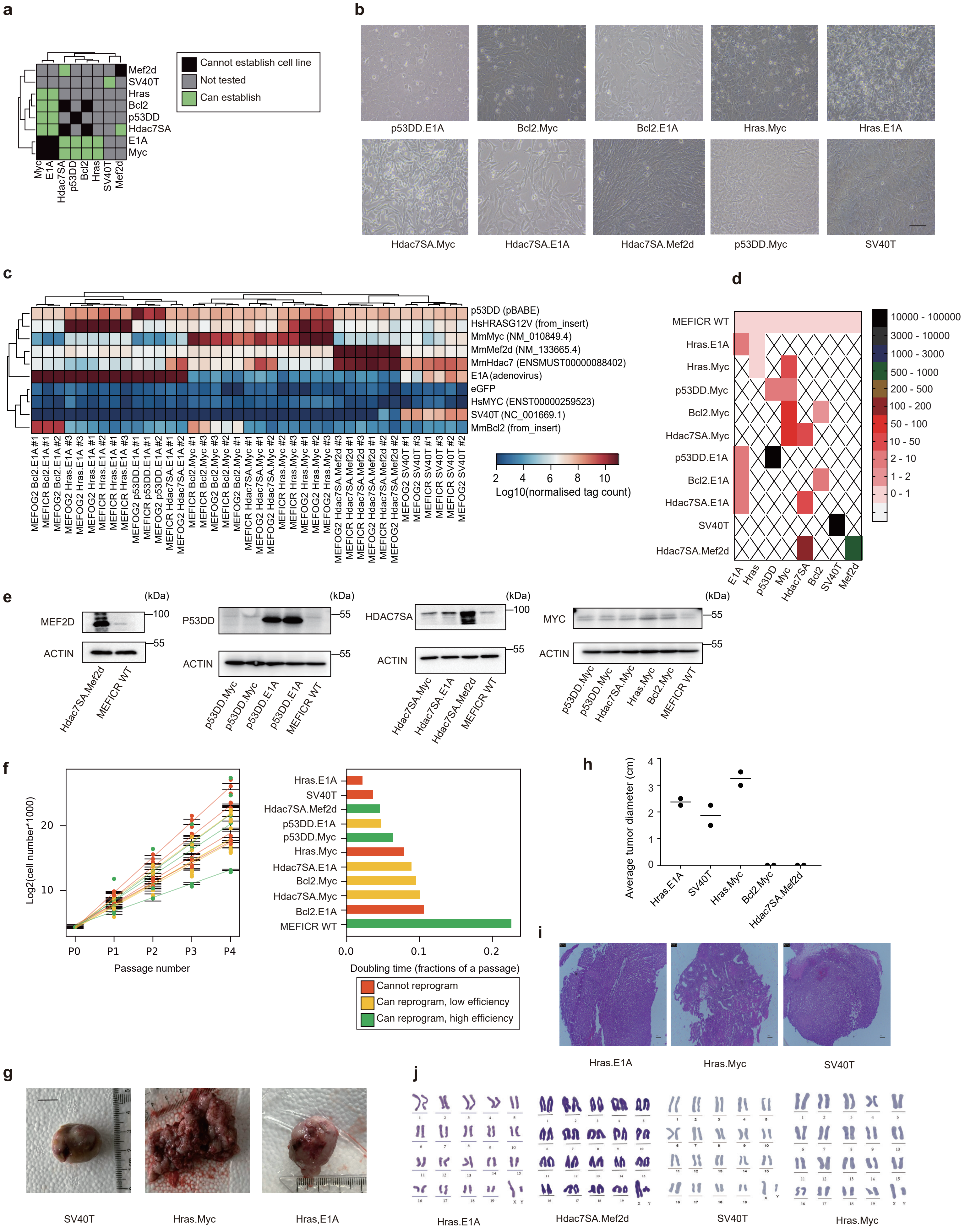


**Supplementary Figure 2**

**Supplementary Fig. 2. Properties of the immortalized MEF lines.**

- **a** Heatmap showing the combinations of transgenes used to establish transformed MEF lines.
- **b** Bright field views of the ICR and OG2-derived MEF lines. Scale bar = 100 $\mu$m.
- **c** Heatmap of the expression for the transgenes used in this study in the OG2 and ICR-derived MEF lines.
- **d** RT-qPCR for the indicated transgenes. Data is the mean of two biological replicates. X indicates not performed.
- **e** Western blot for the indicated transgenes in the indicated cell lines. Beta-Actin was used as a control.
- **f** Growth curves of the immortalized MEFs (left chert), and estimated doubling time (right chart). Dots are the mean from three biological replicates. Bars indicate the standard deviation.
- **g** Representative tumor images from MEF-derived growths in nude mice, images. Scale bar = 1cm
- **h** Chart of tumor sizes for the indicated transformed lines. The middle bar represents the mean of the two biological replicates.
- **i** Hematoxylin and eosin staining of cross sections of tumors. Scale bar = 100 $\mu$m
- **j** Karyotype of the indicated MEF lines.

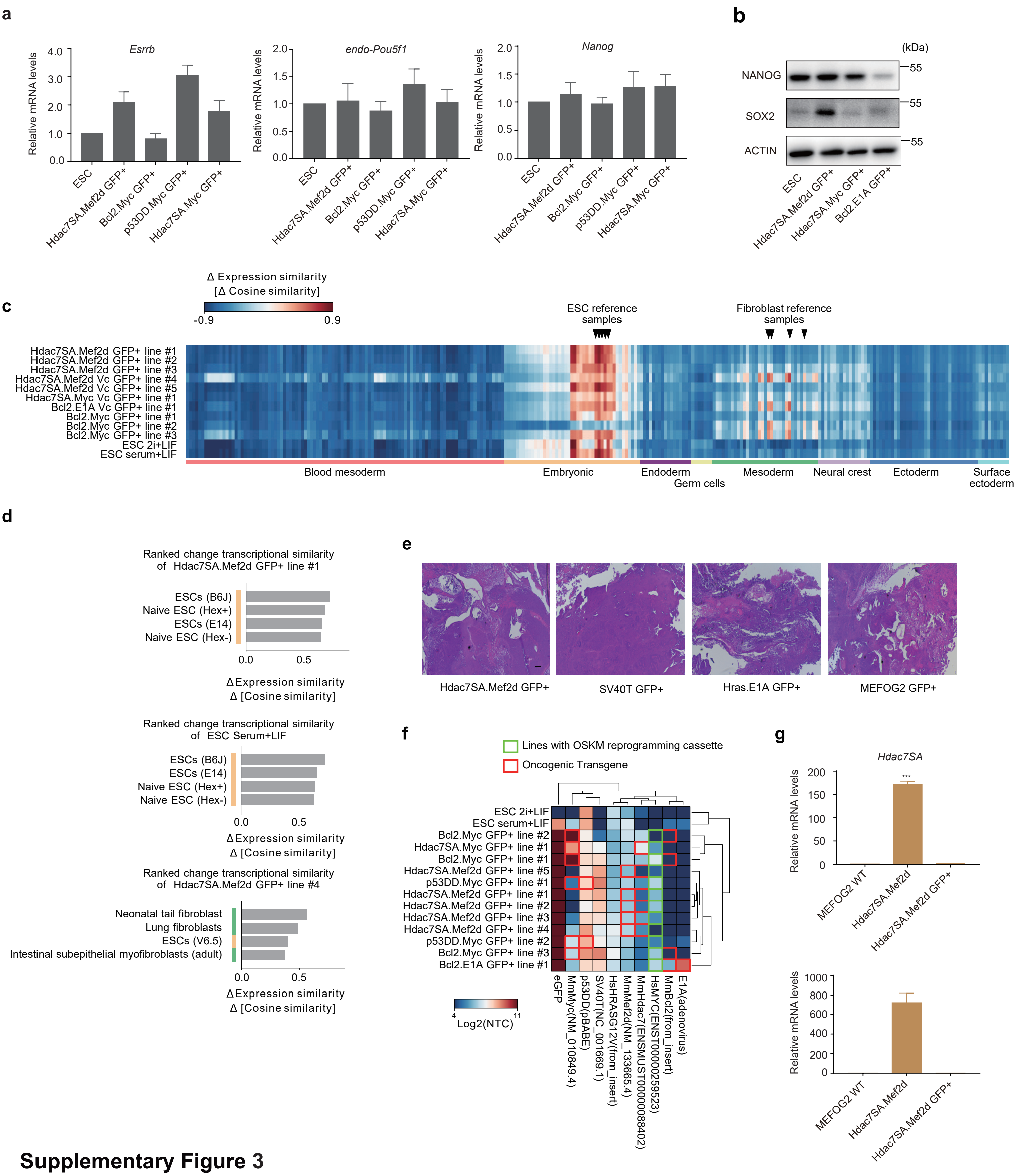


### Supplementary Figure 3

**Supplementary Fig. 3. Reprogramming status of the iPSC lines derived from the immortalized MEFs.**

- **a** RT-qPCR for a selection of pluripotency-related genes in the GFP+ cells at day 15.
- **b** Western blot for SOX2, NANOG and ACTIN (control) in the ESCs and the indicated transformed MEF lines.
- **c** Correlation of the iPSCs versus a large panel of mouse RNA-seq data (Hutchins et al., 2017), using DPre (Steffens et al., 2020).
- **d** Expression similarity as determined by DPre for the Hdac7SA.Mef2d, and ESC lines.
- **e** Representative images Hoechst-stained images from teratomas from the indicated transformed MEF lines.
- **f** RNA-seq data for the transgenes used in this study. (See also Figure S12c). The reprogramming OSKM cassette is a polycistronic vector but only the HsMYC gene is detectable in the RNA-seq data. Note that the p53DD is hard to determine by RNA-seq as the endogenous p53 interferes with the exogenous transgene. Oncogenic transgenes used to generate the MEF lines are indicated in red, and those cells with the OSKM cassette are marked in green.
- **g** RT-qPCR for *Hdac7SA* and *Mef2d* transgenes in the GFP+ iPSC-like lines. The mean of two biological replicates ± s.d. is shown. ***$P < 0.01$.

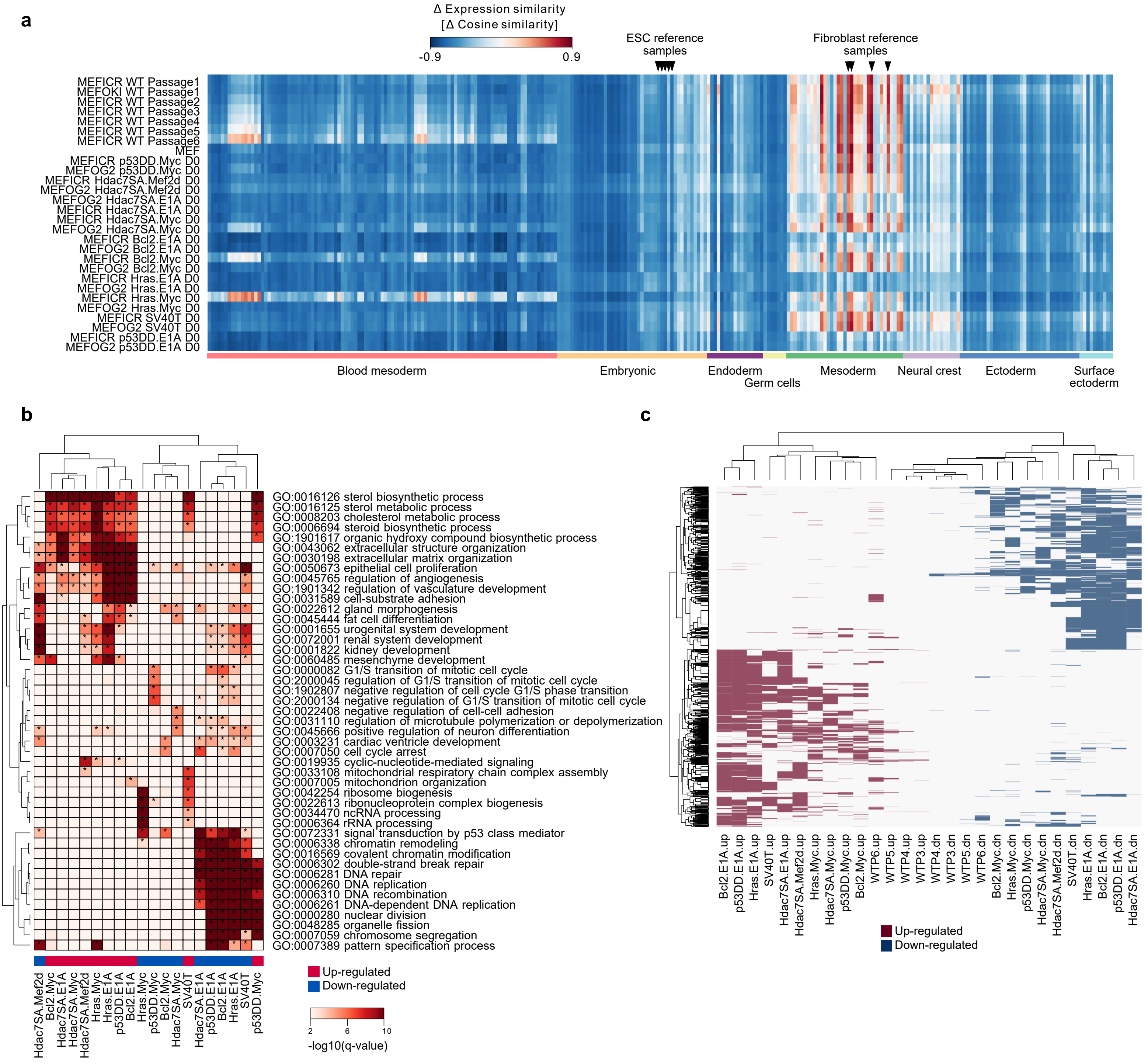


#### Supplementary Figure 4

**Supplementary Fig. 4. Gene expression and epigenetic changes in the transformed immortal MEF lines.**

- **a** No evidence that the MEFs are differentiating, correlations versus the closest samples from (Hutchins et al., 2017) using DPre (Steffens et al., 2020).
- **b** Gene ontology analysis of the DE genes in each MEF line. Up-regulated genes are in the left heatmap and down-regulated genes are in the right heatmap.
- **c** Heatmap of significantly up (red) and down-regulated (blue) genes in the indicated MEF transformed cell lines. Heatmap was generated using any genes that appeared in any four cell lines (either up or down-regulated).

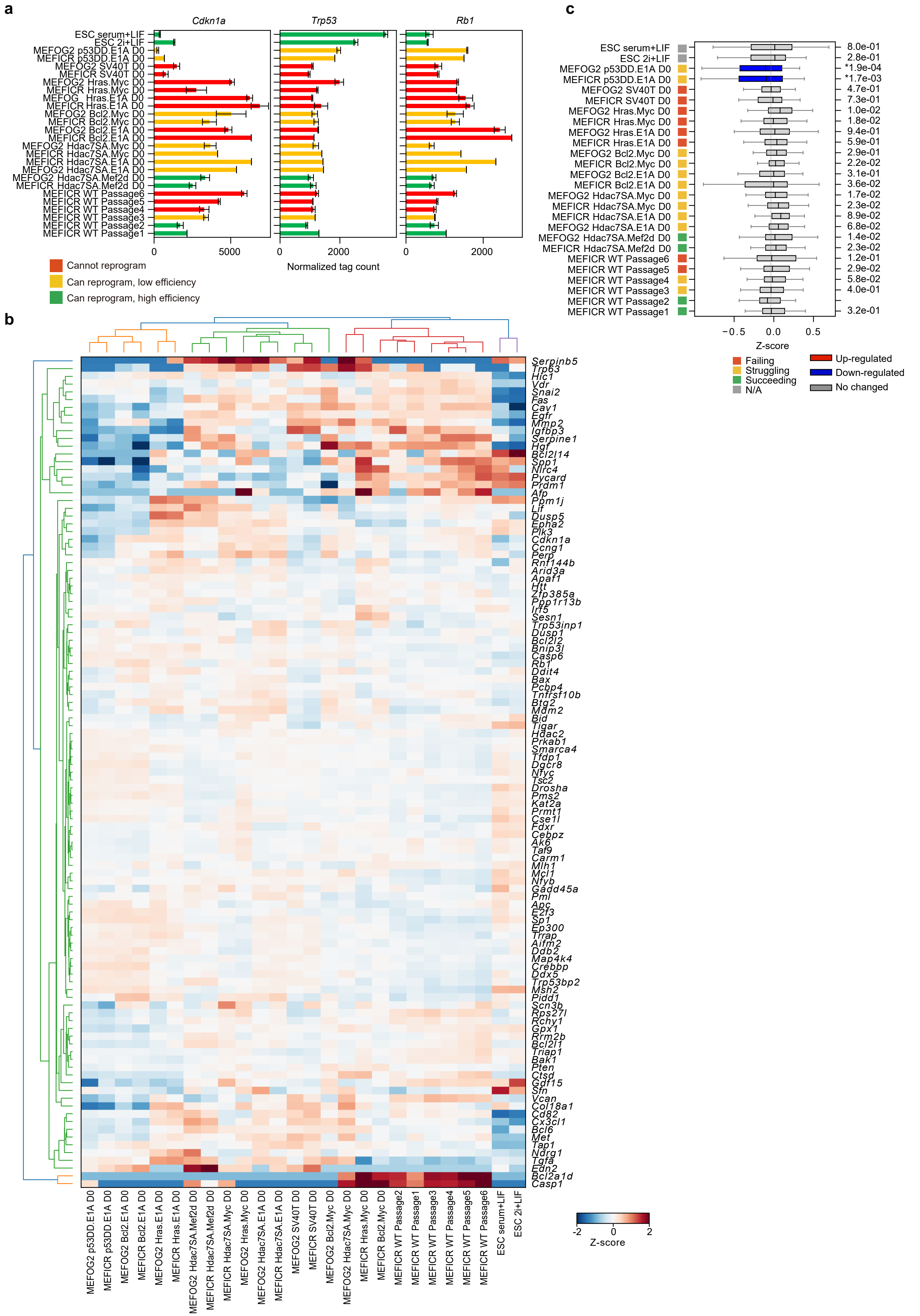


### Supplementary Figure 5

**Supplementary Fig. 5. Known blocks on reprogramming are not responsible for the inability of transformed MEFs to reprogram**

- **a** Bar chart of the expression of *Tp53*, *Cdkn1a* and *Rb1* expression in transformed MEF lines and the first 6 passages of wildtype MEFs
- **b** Z-score heatmap of p53 target genes (taken from PID_P53_DOWNSTREAM_PATHWAY MSIGDB_C2)
- **c** Box plots of the genes in **panel b**, significance is versus MEFICR WT Passage 2 samples, calculated from a two-sided Welch's t-test. Blue indicates significantly down-regulated.

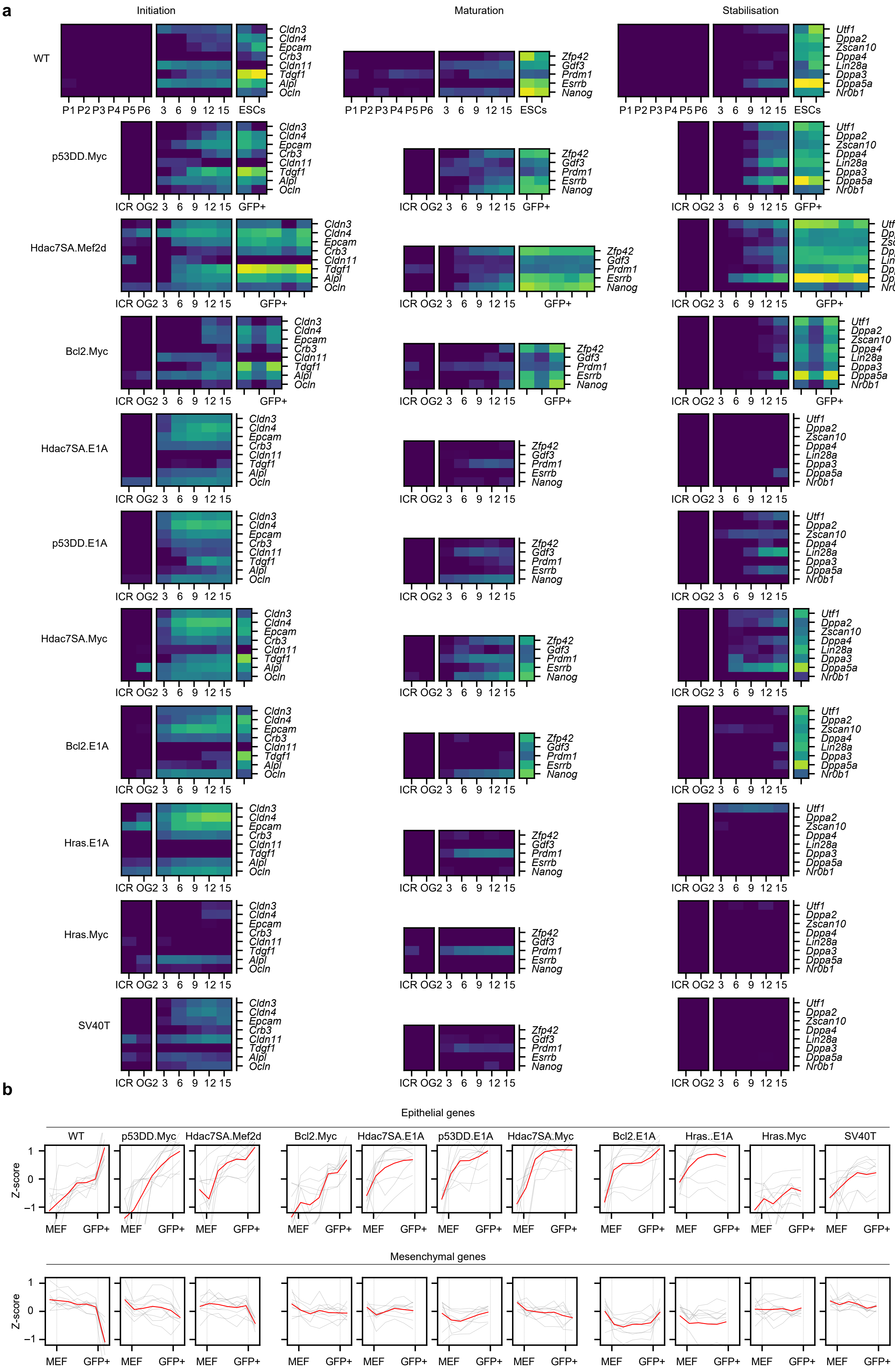


**Supplementary Figure 6**

**Supplementary Fig. 6. Transformed cells have disrupted phases of reprogramming**

**a** Heatmaps showing the expression of initiation (left panel), maturation (middle panel) and stabilization (right panel) genes (Samavarchi-Tehrani et al., 2010), at the indicated time points (days) or in ICR/OG2 MEFs, ESCs, and GFP+ cell lines.

**b** Line charts showing the expression of mesenchymal and epithelial genes at the indicated time points or in MEFs and GFP+ cells. The red line is the mean of all genes, the grey lines are the individual genes.

**a**

| Drug name | Function | Effect |
|---|---|---|
| PD0325901 | ERK/MEK inhibitor | Promote |
| CHIR99021 | GSK inhibitor | Promote |
| Y23637 | ROCK inhibitor | Promote |
| BIX-01294 | Methyltransferase G9a inhibitor | Promote |
| TSA | HDAC inhibitor | Promote |
| Rapamycin | mTOR inhibitor | No effect |
| DZNep | Methyltransferase inhibitor | No effect |
| MS023 | Methyltransferase Pan-PRMT inhibitor | No effect |
| OICR9429 | Methyltransferase WDR-family inhibitor | No effect |
| Chaetocin | Methyltransferase Suv39h1/2 inhibitor | No effect |
| UNC0379 | Methyltransferase SETD8 inhibitor | No effect |
| CPTH2 | HAT inhibitor | No effect |
| Garcinol | HAT inhibitor | No effect |
| Sirtinol | Deacetylation SIRT1/2 inhibitor | No effect |
| JQ1 | BRD4 inhibitor | No effect |
| Bromosporine | Bromodomain Pan-BR inhibitor | No effect |
| AZD6738 | Ras inhibitor | No effect |
| BMH-21 | DNA/RNA synthesis inhibitor | No effect |
| CX5461 | DNA/RNA synthesis inhibitor | No effect |
| BX-795 | PDK1 inhibitor | No effect |
| GW843682X | Cell cycle inhibitor | No effect |
| BX-912 | PDK1 inhibitor | No effect |
| 10058-F4 | Cell cycle inhibitor | No effect |
| ICP-203 | Cell cycle inhibitor | No effect |
| Flavopiridol | CDK inhibitor | No effect |

**b**

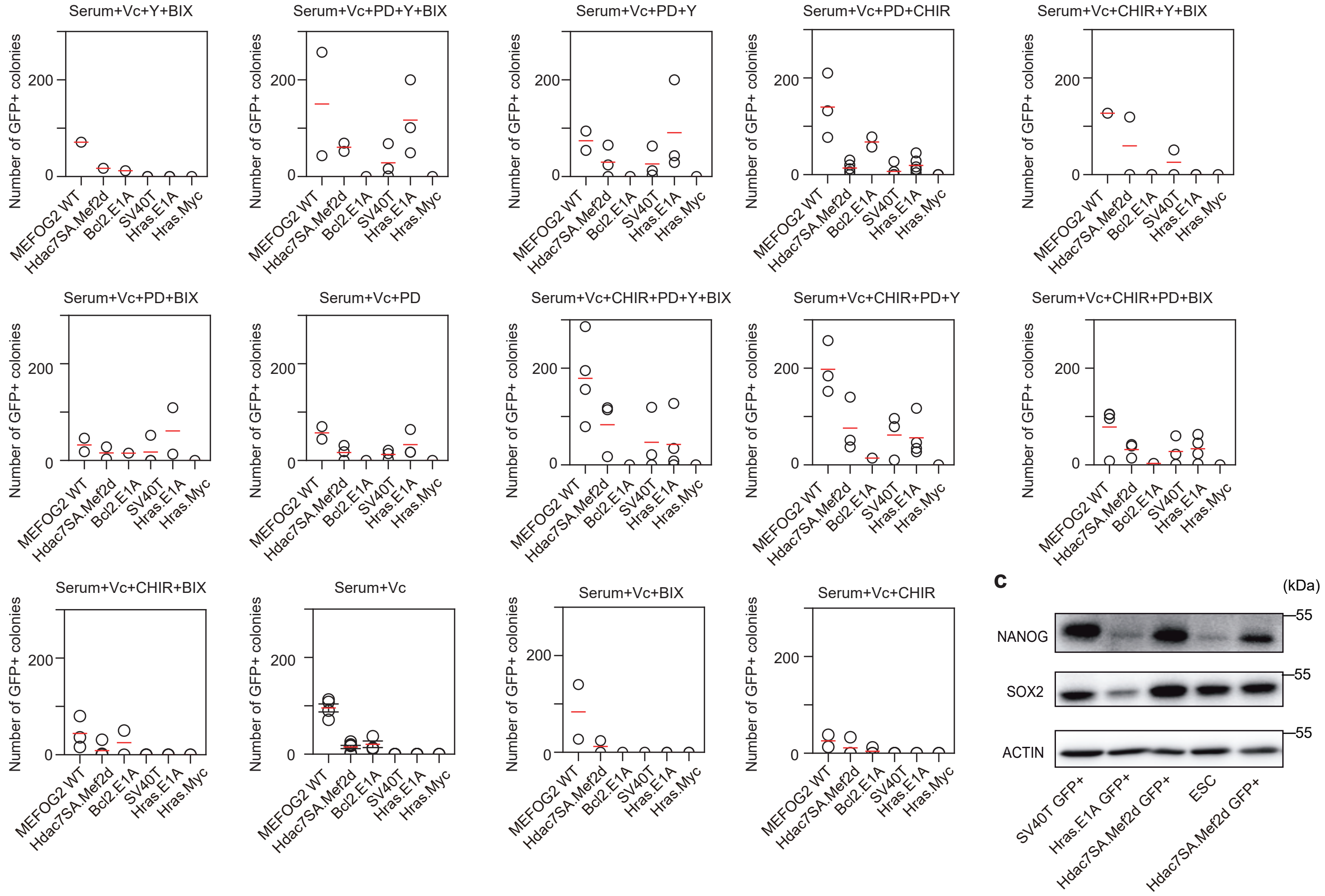


### Supplementary Figure 7

**Supplementary Fig. 7. Development of the inhibitor cocktail to convert reprogramming-incapable cells to -capable.**

- **a** A summary of this inhibitors used in this study.
- **b** GFP+ colony counts from reprogramming experiments for the indicated inhibitor cocktail combinations. Data is from a varied number of biological replicates per transformed cell line and inhibitor treatment. Most experiments were performed twice for two biological replicates (Serum+Vc+PD+Y+BIX, Serum+Vc+PD+Y, Serum+Vc+CHIR+Y+BIX, Serum+Vc+PD+BIX, Serum+Vc+PD, Serum+Vc+BIX, Serum+Vc+CHIR) or in triplicate (Serum+Vc+PD+CHIR, Serum+Vc+PD+CHIR+Y, Serum+Vc+PD+CHIR+BIX, Serum+Vc+CHIR+BIX), or in quadruplicate (Serum+Vc+PD+CHIR+Y+BIX, Serum+Vc+PD+CHIR+BIX, serum+Vc). One experiment was not replicated (Serum+Y+BIX).
- **c** Western blot for NANOG, SOX2 and ACTB as a loading control in the indicated GFP+ lines derived from the indicated WT or transformed MEFs.

**a**

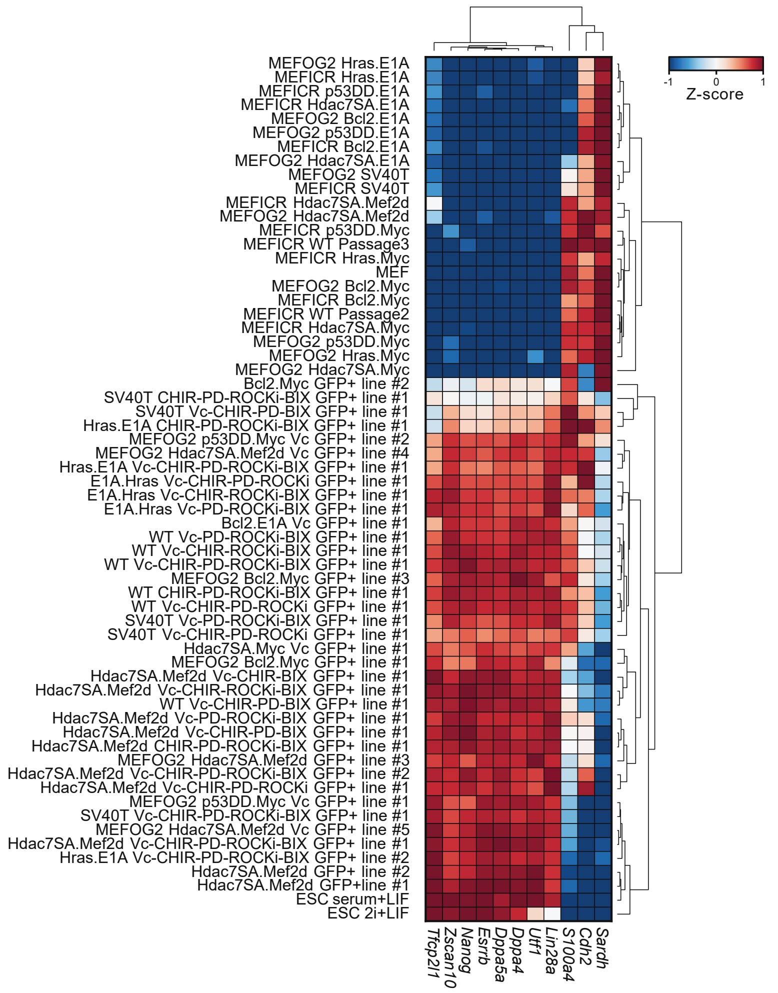


**b**

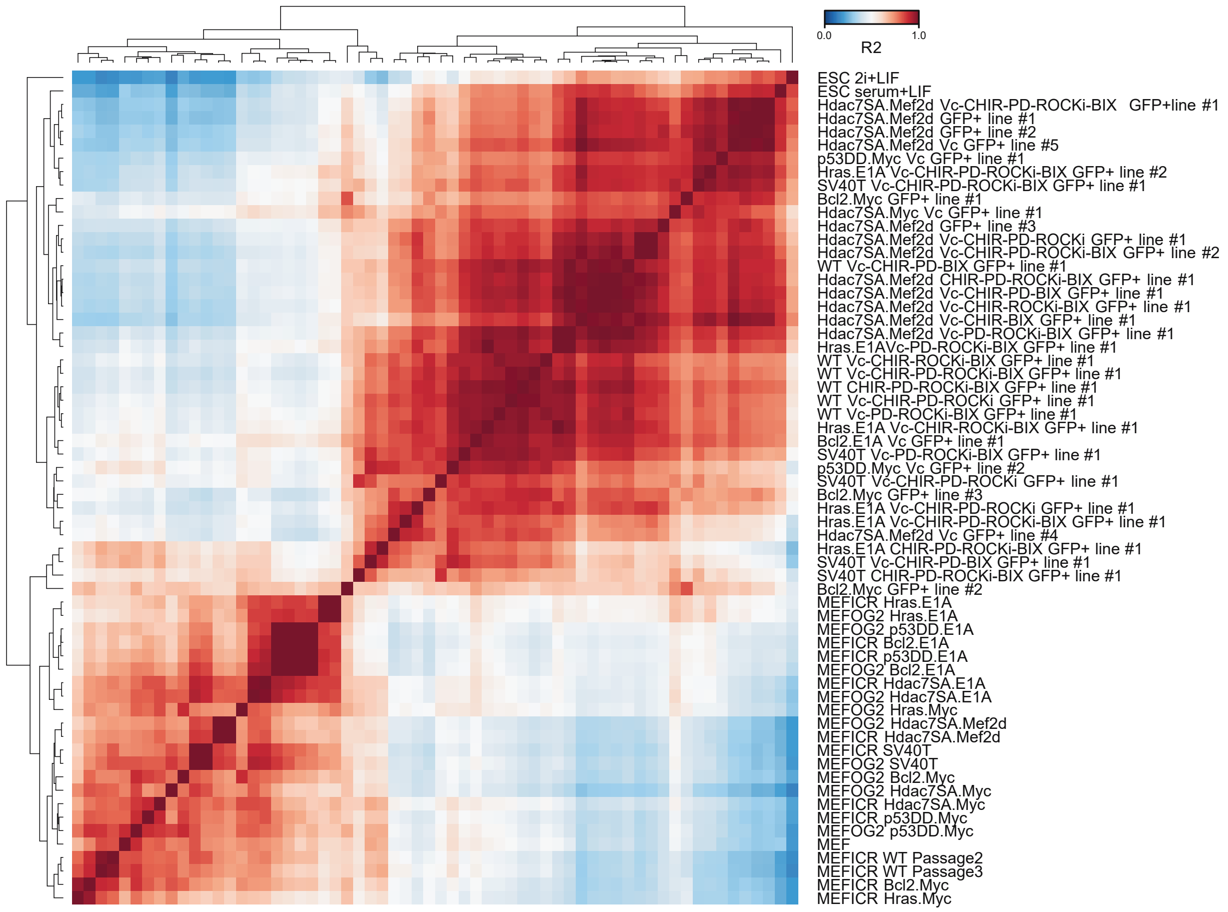


#### Supplementary Figure 8

**Supplementary Fig. 8. Reprogrammed cell lines closely resemble wildtype ESCs**

**a** Heatmap of the expression of selected pluripotency and somatic cell factors in the WT MEFs, transformed MEF cells lines, WT ESCs and the GFP+ sorted cell lines derived from the transformed MEFs. Chemical conditions used in the reprogramming process are indicated for each sample. If not stated then reprogramming was performed in serum+Vitamin C.

**b** Cross-correlation ($R^2$) of the RNA-seq data for the WT MEFs, transformed MEF cells lines, WT ESCs and the GFP+ sorted cell lines derived from the transformed MEFs.

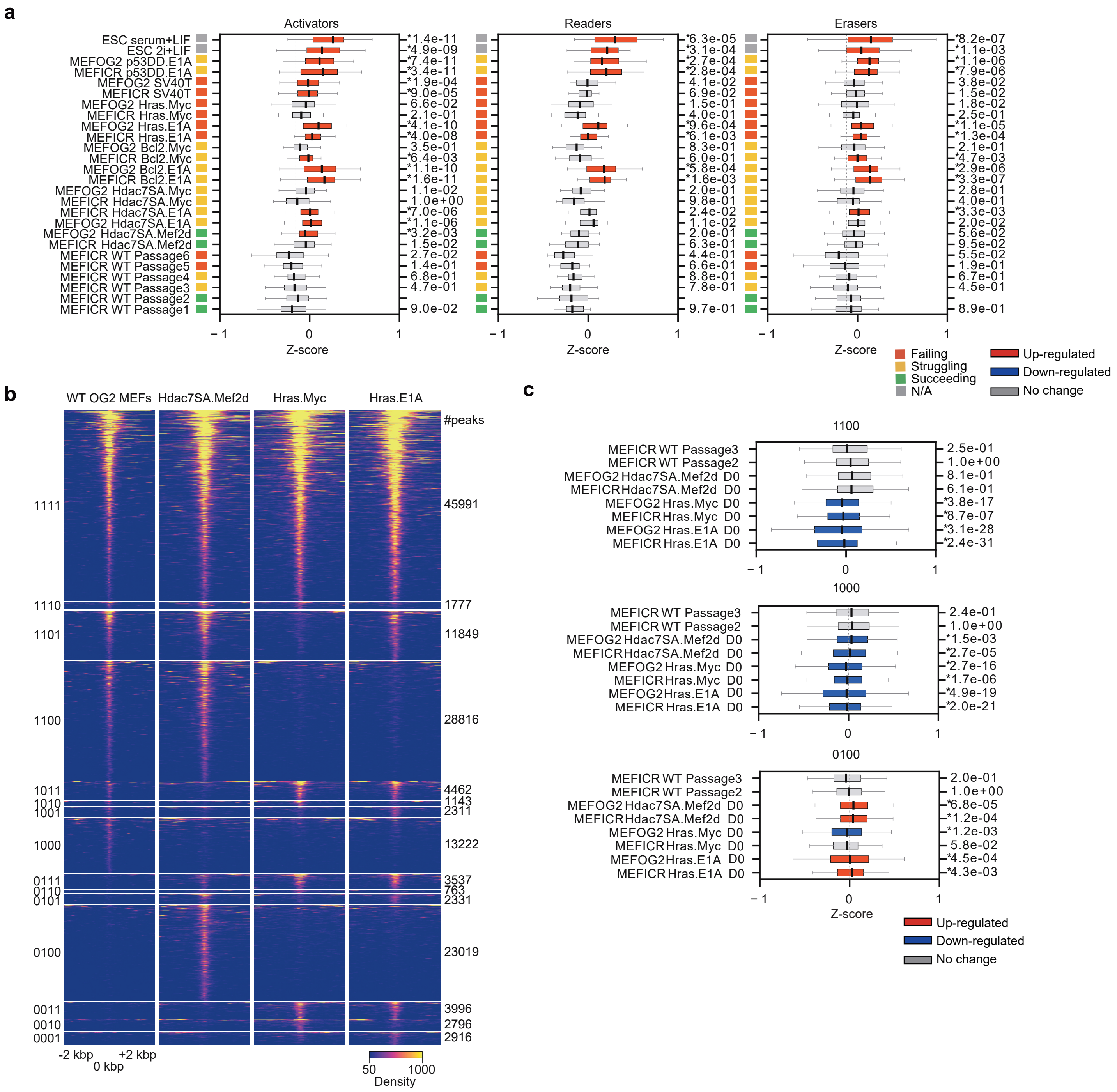


#### Supplementary Figure 9

**Supplementary Fig. 9. Epigenetic control of reprogramming-capable and -incapable lines**

- **a** Box plots of the expression of epigenetic factors in MEFs. Epigenetic factors were downloaded from the Epifactors database (Medvedeva et al., 2015), and allocated to one of three categories: Activator, Reader or Eraser. The Z-scores of expression is indicated. Significance is from a Welch's test with MEFICR Passage2 compared to all other samples. Changes were considered significant if the p-value was < 0.01.
- **b** Heatmap comparing the ATAC-seq data in WT MEFs to the ATAC-seq in the indicated transformed cell lines, for the indicated binary classifications of open (1) or not open (0) shown on the left side of the heatmap. The plot is centered on the peak summit and shows the flanking 1 kbp. Normalized tag counts are shown for the indicated samples.
- **c** Boxplots of the expression of all genes within 2 kbp of a set of loci that are specifically opened or closed in the indicated lines. Three groups of peaks are shown, 1100 (open in WT MEFs and Hdac7SA.Mef2d, closed in the other lines), 1000 (only open in WT MEFs), and 0100 (specifically open in the Hdac7SA.Mef2d).

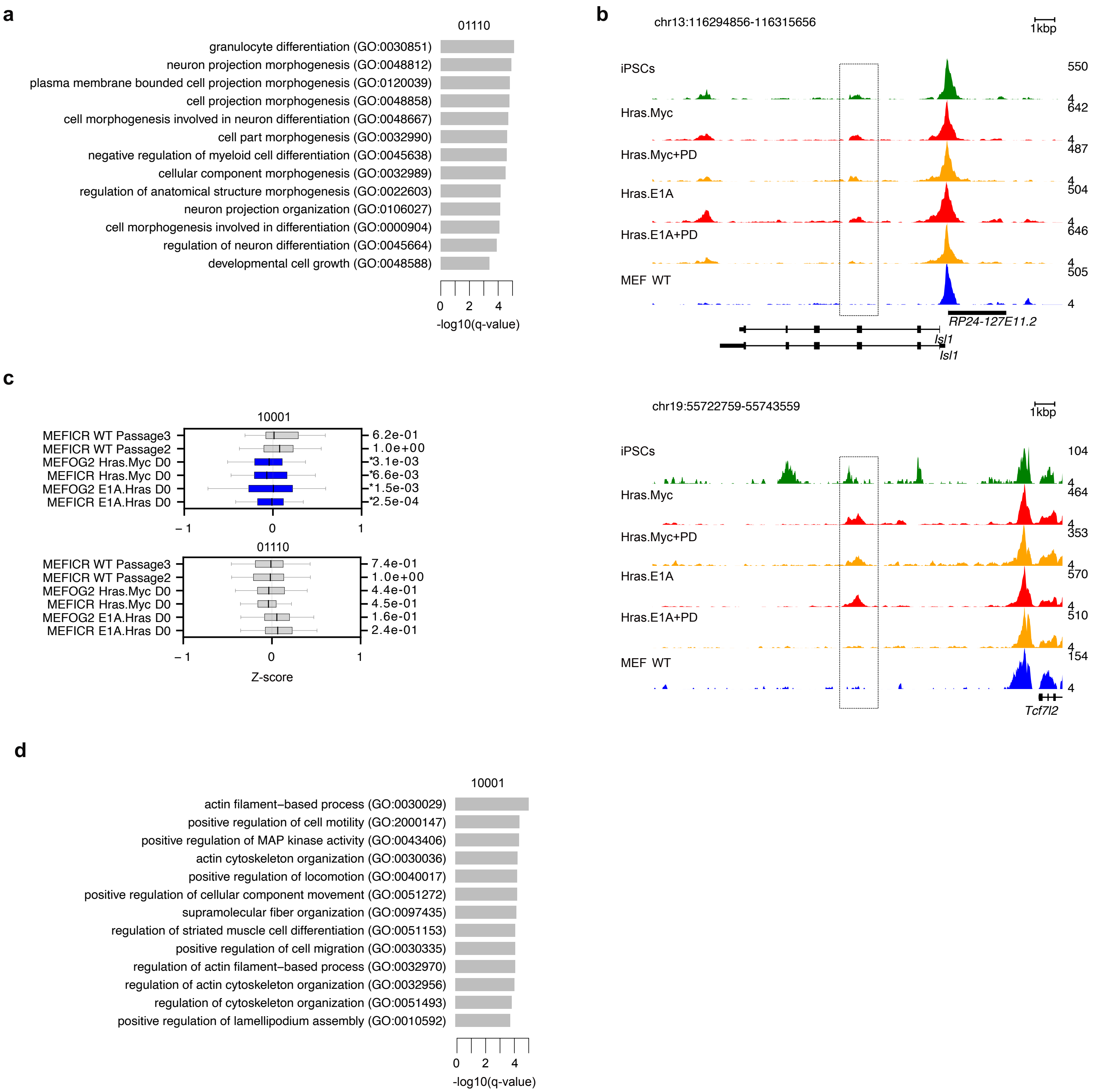


#### Supplementary Figure 10

**Supplementary Fig. 10. Chromatin at developmental enhancers is suppressed by MEK inhibition**

- **a** GO analysis of the genes within 5000 bp of a chromatin locus in the 01110 group.
- **b** Genome views of two developmental genes, Isl1 and Tcf7l2.
- **c** Boxplots showing the gene expression of the genes within 5000 bp of a chromatin locus in the 01110 or 10001 groups. Significance is from a two-sided Welch's t-test.
- **d** GO analysis of the genes within 5000 bp of a chromatin locus in the 10001 group.
